# Biophysical modeling identifies an optimal hybrid amoeboid-mesenchymal mechanism for maximal T cell migration speeds

**DOI:** 10.1101/2023.10.29.564655

**Authors:** Roberto Alonso-Matilla, Diego I. Pedro, Alfonso Pepe, Jose Serrano-Velez, Michael Dunne, Duy T. Nguyen, W. Gregory Sawyer, Paolo P. Provenzano, David J. Odde

## Abstract

Despite recent advances in understanding cell migration mechanics, the principles governing rapid T cell movement remain unclear. Efficient migration is critical for antitumoral T cells to locate and eliminate cancer cells. To investigate the upper limits of cell speed, we developed a hybrid stochastic-mean field model of bleb-based cell motility. Our model suggests that cell-matrix adhesion-free bleb-based migration is highly inefficient, challenging the feasibility of cell swimming/adhesion-independent migration as a primary fast motility mode. Instead, we show that T cells can achieve rapid migration by combining bleb formation with adhesion-based forces. Supporting our predictions, our three-dimensional gel experiments confirm that T cells migrate significantly faster under adherent conditions than in adhesion-free environments. These findings highlight the mechanical constraints of T cell motility and suggest that modifying tissue adhesion properties in a controlled manner could enhance immune cell infiltration into tumors. Our computational and experimental work provides insights for optimizing T cell-based immunotherapies. While antifibrotic treatments could alter the tumor microenvironment, indiscriminate reduction of adhesion may not be ideal for T cell infiltration and motility, highlighting the need for targeted antifibrotic strategies.

---

Cell migration is a complex multistep process that is critical for *in vivo* processes such as morphogenesis^1^, maintaining tissue health and homeostasis^2^, and driving diseases such as cancer, including cancer cell dissemination during metastasis^3^ and immune cell migration within tumors^4^. Single cells migrating in three-dimensional environments use two dominant modes of migration: amoeboid and mesenchymal^5^. The amoeboid phenotype is characterized by a rounded cellular morphology, high cortical contractility, low cell-matrix adhesions and narrow/tight extracellular spaces, and it is commonly associated with the formation of blebs^6-13^, which are pressure driven hemi-spherical cellular membrane protrusions that are initiated by either a local membrane-cortex detachment or local cortex rupture^14-16^. This amoeboid blebby phenotype has been reported under some conditions for transformed cells (e.g. melanoma, breast and squamous carcinoma cells *in vivo;*^17-20^), and is frequently utilized by T cells^21-23^. The mesenchymal-like phenotype is frequently displayed by fibroblasts and multiple cancer types, including carcinomas, fibrosarcomas and glioblastoma^24,25^. It is characterized by an elongated morphology, strong cell-matrix adhesion, and the formation of actin-rich leading-edge structures such as spike-like filopodia or broad flattened lamellipodia^10,12,24,26^. Cells navigate through extremely complex microenvironments, such as tight porous viscoelastic tissues or narrow vascular pores. As a result, many cancer cells^5,12,17,27-31^ and immune cells, including T cells^21^, can extend both types of cellular protrusions, and adapt to different mechanical and chemical microenvironments by transitioning between mesenchymal and amoeboid migration phenotypes.

T cell-based cancer therapies offer a promising approach in the fight against cancer^32^. However, successful T cell-based therapies have been limited mainly to hematologic cancers. Solid tumors create a highly immunosuppressive and fibrotic microenvironment that hinders the physical infiltration and activity of cytotoxic T cells, thus enabling tumor growth and metastasis^33,34^. Despite the significant progress in understanding bleb formation in recent years^21,35,36^, it remains largely unknown how bleb-producing T cells exchange momentum with their surroundings to navigate through challenging mechanical and chemical microenvironments such as tumors.

Adhesion-dependent migration, described and mathematically modeled using a motor-clutch framework^25^, has long been thought to be required for cell motility^37^, but some recent findings have challenged this view^11,26,38-51^. Following the seminal paper by Lämmermann et al.^38^, which demonstrated that leukocyte migration *in vitro* and *in vivo* is independent of various integrin heterodimers, there has been an ongoing debate about whether the rapid migration of immune cells, particularly those exhibiting an amoeboid phenotype, requires adhesion-based forces^40,44,48,51^. The question of whether immune cells can swim through tissues autonomously or if they depend on other adhesion molecules besides integrins (e.g. CD44) for migration remains unresolved. Notably, T cells are capable of achieving exceptionally high migration speeds, with *in-vivo* motility coefficients of approximately 10 µm^2^ /min21, a phenomenon that remains poorly understood and raises questions about potential limits imposed in standard adhesion-based motor-clutch migration mechanisms by diffusion-limited F-actin polymerization rates. Additionally, while adhesion-independent models have been proposed, a recent study^52^ has shown that T cells can generate measurable contractile forces on compliant substrates, suggesting that traction forces and substrate interactions may still play an important role during T cell migration.

Most animal cells have a dense, thin actomyosin layer underneath the plasma membrane that produces cortical tension mediated forces that are transmitted to the membrane by membrane-cortex linker proteins, such as Ezrin, Radixin and Moesin proteins^53,54^. Upon local membrane-cortex cohesion loss, cortical forces no longer get transmitted to the plasma membrane and a bleb nucleates and undergoes rapid expansion reaching bleb sizes of order ∼1 μm. At the bleb nucleation site, the local hydrostatic pressure presumably drops, and cytoplasmic material flows from the center of the cell to the low-pressure bleb site^55,56^. The bleb cycle ends by the self-assembly of new cortex underneath the bleb membrane, which drives bleb retraction^14,57^. Yet, it is not obvious whether this form of cell motility requires cell-matrix adhesion interactions, nonspecific cell-matrix friction interactions or if the cell can achieve a net displacement after each bleb cycle in adhesion-independent conditions. Hence, a combination of biophysical modeling and cutting-edge experimental approaches is needed to elucidate T cell migration mechanisms and establish design criteria for next-generation immune engineering strategies to enhance T cell infiltration, migration, and sampling within the tumor mass.

Bleb nucleation^58-63^ and bleb growth^56,64-66^ have been studied theoretically using coarse-grained mechanistic models. In addition, a handful of amoeboid/bleb-based motility biophysical models have also been recently developed, where the intracellular and extracellular spaces were assumed to behave as viscous fluids^49,50,67-79^. Most of these cell migration models assume that plasma membrane and cortex are a single composite material. Two of these studies, however account for the mechanical interactions between the intracellular viscous fluid, the actin cortex and the cell membrane^67,68^. However, the model by Lim et al.^67^ does not satisfy conservation of momentum, since it does not account for the forces exerted by the cortex on the cytoplasm as it moves through the cortex to fill up the expanding bleb. Likewise, the model of Copos and Strychalski^68^ introduces a cortical viscoelastic force that depends on a reference cortical configuration, as well as a steric interaction force with adjacent confining walls. Although the enforced no-penetration fluid flow condition at the boundaries ensures conservation of linear momentum in the fluid, the inclusion of these two forces does not guarantee that the total force exerted by the cell on the medium vanishes (i.e. does not ensure conservation of overall momentum). To address this gap, in the present study, we first develop a momentum-conserving mechanistic model of bleb-based cell swimming in a viscoelastic medium to assess whether bleb-producing cells such as T cells can mechanically swim in the absence of adhesions, identify the key cellular components that regulate the motile capabilities of swimming cells, and quantify the theoretical maximum speed that a cell can achieve using this migration mechanism. Our initial focus on cell swimming is motivated by the persistent question of how T cells navigate through distinct microenvironments, ranging from lymph nodes to extracellular spaces of varying densities. We find that while bleb-based cell swimming is theoretically possible, it does not seem to be sufficient on its own to explain the rapid T cell migration observed experimentally in vivo. Rather, our analysis supports an alternative hybrid model in which blebbing drives rapid protrusion extension, while subsequent weak and transient adhesion stabilizes the newly gained cell position. This proposed migration mechanism is reinforced by T cell experiments in three-dimensional liquid-like solid granular gels, demonstrating that adhesion-based forces are essential for effective migration. Our work provides key insights to mechanically control and potentially maximize migration of blebby cells such as T cells that can contribute to the design of tumor stroma-targeting and cell-specific engineering strategies to enhance the intratumoral migration capabilities of antitumor immune cells.

## RESULTS

### Biophysical model of bleb-based cell swimming

We first aim to investigate the cell migration capabilities of bleb-producing cells in an unbounded viscoelastic medium through the development of a two-dimensional biophysical mechanistic model (see Supplemental Material for a detailed description of the model). The cell is composed of two distinct structures: a cellular membrane and an actomyosin cortex (Fig. 1A). Both are considered Lagrangian structures defined by a set of nodes that are connected to each other and the associated fluids as described below and in more detail in the supplement. As the cell migrates through the environment, both membrane and cortex structures transmit forces on the surrounding medium, generating viscoelastic stresses. At the continuum level, conservation of mass and momentum on the incompressible viscoelastic fluid reads

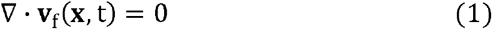

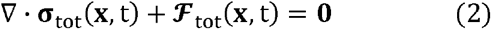

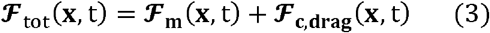

where **v**_**f**_ is the fluid velocity, **σ**_**tot**_ is the total stress, **ℱ**_**tot**_ is the total force density on the fluid and equal to the sum of the membrane force density **ℱ**_**m**_ and cortical force density **ℱ**_**c**,**drag**_, generated in the intracellular medium. The total stress **σ**_**tot**_ = **σ**_**f**_ + **σ**_**p**_ is the sum of two contributions: a purely viscous stress **σ**_**f**_ and a viscoelastic stress **σ**_**p**_:

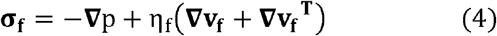

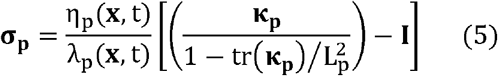

where we have chosen the FENE-P constitutive relation for the stress to model the viscoelastic nature of the intracellular and extracellular spaces^80^. Here, p is the hydrostatic pressure, η_f_ and η_p_ are, respectively, the fluid viscosity and polymer viscosity, λ_p_ is the polymer stress relaxation time, **κ**_**p**_ is the conformation stress tensor and L_p_ is the polymer extensibility parameter, a measure of the maximum polymeric deformation. The time evolution of the conformation stress tensor **κ**_**p**_ reads

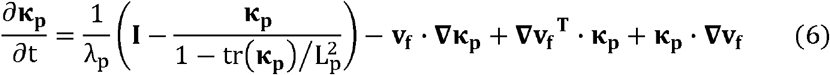

**Figure 1.**
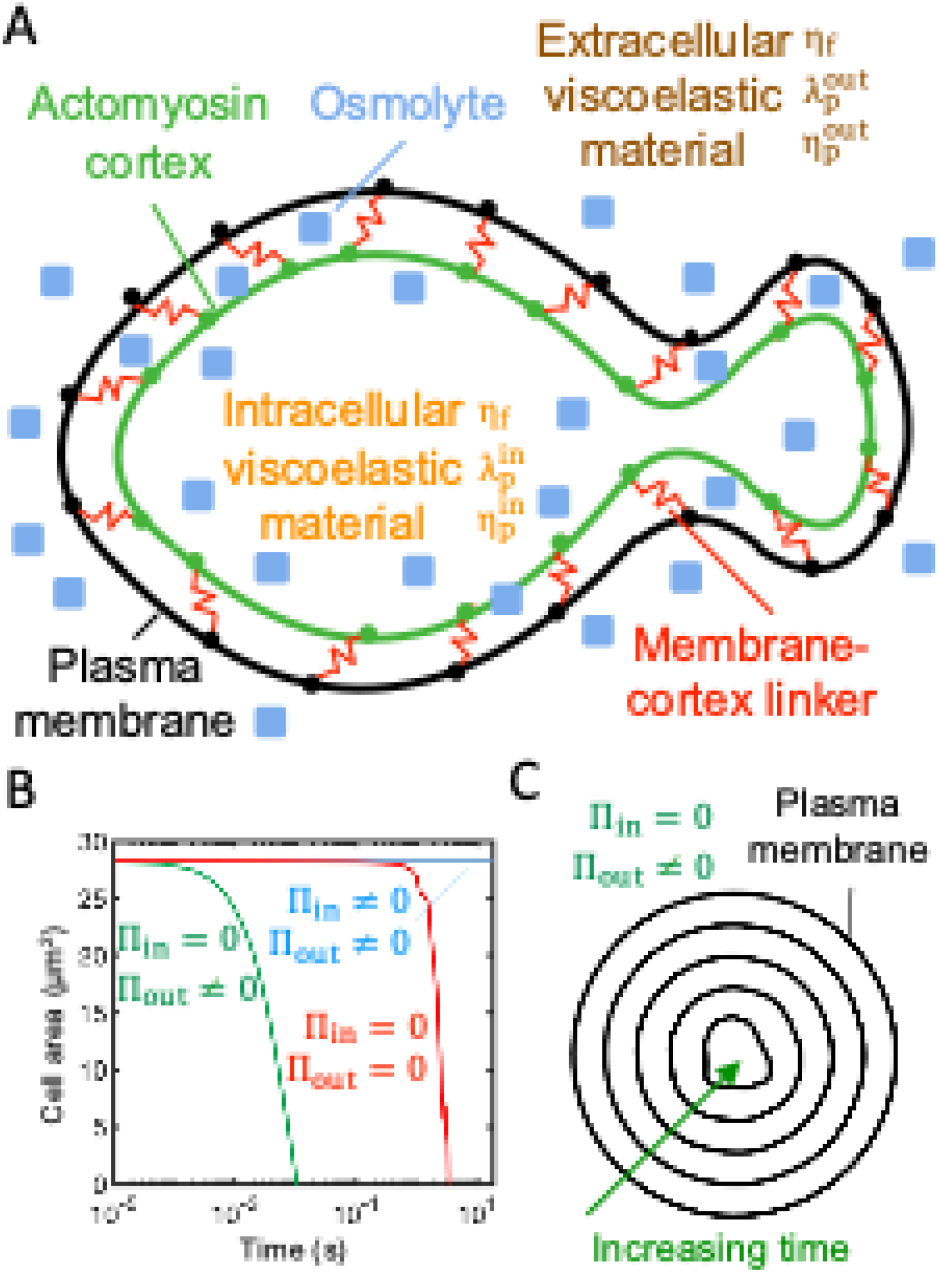
Intracellular osmotic pressure sets a stable cell size by counteracting cell shrinkage mediated by extracellular osmotic pressure and cortical tension. (A) Schematic diagram of the bleb-based cell swimming model. Plasma membrane and actomyosin cortex are mechanically linked by reversible Hookean elastic linkers that stochastically associate and dissociate in a force-independent and -dependent manner, respectively. Intracellular and extracellular spaces are viscoelastic materials with associated fluid viscosity η_f_, polymer viscosity η_p_ and polymer stress relaxation time λ_p_. Osmotic effects are included. (B) Time-evolution of cell area for three different osmotic conditions. Intracellular osmotic pressure: Π_in_, extracellular osmotic pressure: Π_out_ (C) Cell shrinkage due to a reverse osmotic shock (Π_in_ = 0). Thus, inclusion of intracellular osmolytes is necessary to prevent cell collapse due to inward cortical tension and extracellular osmotic forces that tend to force water out of the cell.

The first term on the right-hand side (RHS) of Eq. (6) captures stress relaxation kinetics, the second term captures polymer advection, and the last two terms capture rotation and deformation of the polymeric material.

The cell membrane is an elastic structure subjected to tension, bending and membrane-cortex adhesion forces, and the cortex is considered an actomyosin poroelastic structure that generates myosin-mediated forces that obey a linear force-velocity relationship and create tension in the cortical network^61,81-83^. The actin cortex experiences drag forces as cytoplasmic material flows through the porous actomyosin network. Mass conservation of membrane-cortex linkers, and actin and myosin in the cortex follows simple stochastic association and dissociation kinetics. No-slip between the plasma membrane and viscoelastic fluids is enforced. Furthermore, the model accounts for osmotic effects, which we found were essential for maintenance of cell volume. The intracellular osmotic pressure resists cortex and extracellular osmotic inward forces, all forces combined set cell size and shape. A hypothetical complete depletion of intracellular osmolyte would cause a sudden cell shrinkage within a few milliseconds as the cortex would otherwise squeeze out all the intracellular water (Figs. 1B, C). The model equations are solved using an immersed boundary method^84^. The specific model parameter values used are summarized in Table S2, unless stated otherwise in the text/figure captions, and the code used for the T cell migration biophysical model is available in reference^85^. Additional details can be found in the Supplemental Information.

### Single bleb formation, expansion and retraction allows cells to swim in viscoelastic environments

Using the above framework, we proceeded to study cell dynamics during a single bleb cycle by enforcing local membrane-cortex detachment. Membrane-cortex uncoupling is followed by a drop in local intracellular hydrostatic pressure and cytoplasmic viscoelastic fluid material flows through the detached actomyosin meshwork from the center of the cell to the low-pressure region, mediating bleb expansion (Figs. 2A, B, S1A, S1B, and movie S1). The poroelastic permeable cortex does not sustain pressure forces and moves retrogradely towards the cell center^86,87^, with actomyosin contractile inward forces balanced by cytoplasm-mediated drag forces, which arise as cytoplasmic material flows through the porous actomyosin network (see Eq. S16). Notice that, in the extreme case where the cortex-cytoplasm drag coefficient is extremely high, the actin meshwork moves along with the cytoplasm, preventing the cytoplasm from filling up the bleb. Eventually, cortex components that detach from the membrane disappear via turnover kinetics (i.e. first-order disassembly without concomitant assembly; see Eqs. S20-21). The bleb cycle ends by recruitment of new cortex constituents underneath the bleb membrane, which nucleates new actin filaments that recruit myosin motors that transmit inward forces to the plasma membrane driving bleb retraction and cytoplasm from the bleb region to the center of the cell in a low intracellular hydrostatic pressure gradient phase (Figs. 2A, 2B and S2). We observe that the bleb area at the end of the cycle is primarily limited due to the increase in the local hydrostatic pressure from cortex reformation rather than membrane tension (Fig. S2). We observe that the plasma membrane follows similar trajectories during bleb expansion and bleb retraction. A forward movement of cytoplasmic material translocates the center of mass of the cell forward in the direction of the bleb during bleb growth, and a rearward movement of cytoplasmic material translocates the center of mass of the cell in the opposite direction during bleb retraction (Figs. 2C–E, S3 and S4). According to Purcell’s scallop theorem, cell shape-based swimming of microscale biological cells in a Newtonian fluid at low Reynolds number requires non-reciprocal body deformations to generate locomotion^88^. Accordingly, we observe that net cell displacements at the end of the bleb cycle are minimal in Newtonian environments (Fig. S2). This result is in contrast with the model by Lim et al., which shows that individual blebbing events lead to significant net cell displacements. This discrepancy likely arises from a violation of momentum conservation in their model. However, reciprocal motion can lead to net cell motion in viscoelastic environments, as observed experimentally for single-hinge reciprocal microswimmers moving through shear-thickening and shear-thinning fluids^89^. For physiologically relevant model parameters, cell displacements are larger during bleb expansion compared to those during bleb retraction due to the rapid and slow rates of bleb expansion and retraction phases, respectively. The cell shears its environment at distinct rates during both phases, resulting in significant net cell displacements (∼0.4 µm) during a single bleb cycle of ∼2.5 seconds (Figs. 2C–E), for a mean speed of ∼9.6 µm/min. We can estimate the 3D effective cell motility coefficient as 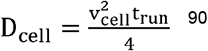, where cells run at an effective velocity given by v_cell_ d_run_ /t_run_ during an effective runtime t_run_. Given that reported blebbing frequencies are on the order of 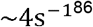, we assume that the time needed for the cell to translocate during each bleb cycle is the limiting factor in each cell run; thus t_run_ ∼2.5s (Figs. 6A, B). Under these assumptions, 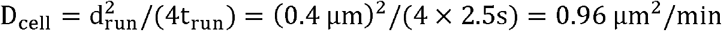. This value is an order of magnitude smaller than the *in-vivo* motility coefficient of bleb-producing cells (∼10 µm /min 21), suggesting that the adhesion-free bleb-based cell migration mechanism is unlikely to be the mode of migration for T cells navigating through tissues. For non-physiological fast cortex turnover times, the timescales of bleb expansion and retraction are similar, preventing cells from achieving net cell motion during a single bleb cycle (Fig. S4). Overall, our results indicate that net cell displacements can be achieved by bleb-based cell swimming in viscoelastic environments, but only at rates insufficient to explain rapid T cell motility *in vivo*.

**Figure 2.**
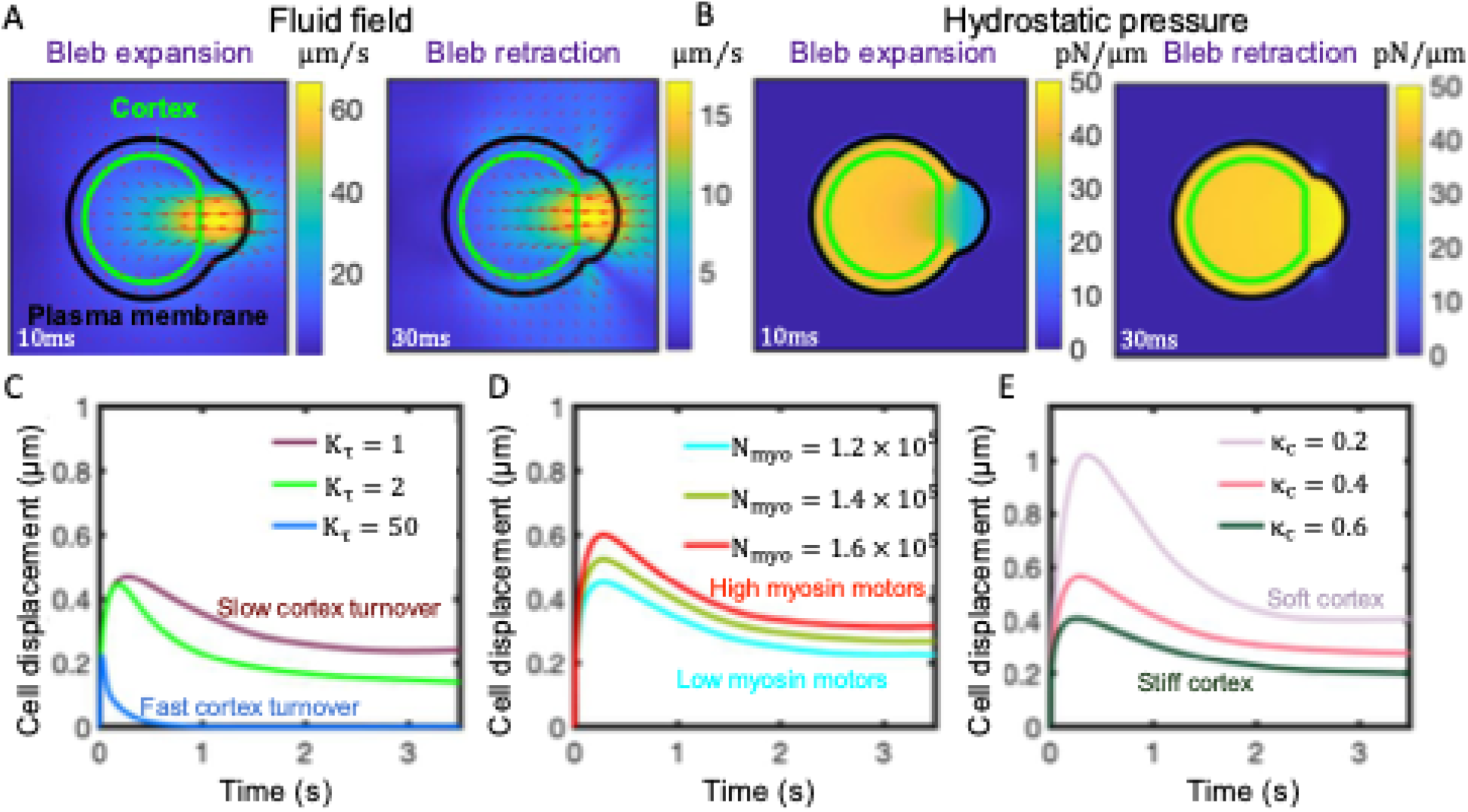
Cells can achieve moderate net cell displacements during a single bleb cycle in viscoelastic environments without adhesion-based forces, due to the differential shear rates associated with bleb growth and retraction phases. (A,B) Snapshots of the magnitude of the fluid velocity and velocity vectors (A), and hydrostatic pressure (B) acquired at two time-points during early bleb expansion and early retraction. K_τ_= 100 for panels (A) and (B). (C E) Time-evolution of the cell centroid for three different values of the cortex turnover parameter K_τ_ (C), for three different values of the number of myosin molecules inside the cell N_myo_ (D) and three different values of the cortical stiffness per unit of actin parameter κ_c_ (E). Greater cell displacements are achieved for slower cortical turnover kinetics, and a more contractile and softer cortex. In all panels, membrane-cortex linkers are broken by hand locally in a small region at time t = 0, mimicking a local downregulation of membrane-cortex linkers due to a chemical signal or force. Polymeric model parameter values: 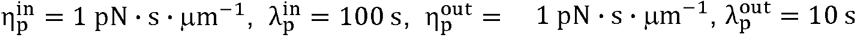.

### High intracellular hydrostatic pressure gradients and a softer cortex favor larger cell displacements during a bleb cycle

Because of uncertainty about our base parameter values, we explored how varying cellular conditions influenced cell displacements during an individual bleb cycle. For a given membrane-cortex detachment size, more contractile and/or softer actomyosin cortices favor larger cell displacements during bleb expansion (Figs. 2D and 2E), resulting in greater overall cell displacements at the end of the bleb cycle. High cortical squeezing, mediated by a higher number of myosin motors in the cell, causes larger intracellular hydrostatic pressure gradients after membrane-cortex dissociation. This drives stronger cytoplasmic flows and results in larger bleb sizes and cell displacements during an individual bleb cycle (Fig. 2D). A softer cortex gives rise to stronger intracellular flows (Fig. S1A), and delays both intracellular hydrostatic pressure equilibration (Fig. S1B) and the end of bleb expansion (Fig. 2E). During this expansion phase, active stresses are concentrated at the cell’s rear and center, away from the actomyosin-free bleb region. The cortex, being a highly crosslinked actin network, allows these active stresses to propagate throughout the cell^91^. As new cortex is recruited beneath the plasma membrane bleb, it experiences retrograde pulling forces from the highly contractile regions away from the bleb, transmitting these forces to the plasma membrane at the leading edge. In cells with a softer cortex, this stress propagation is diminished, resulting in delayed bleb retraction and larger cell displacements at the end of a bleb cycle (Fig. 2E). Interestingly, bleb expansion rates obtained from Fig. 2 (∼5 − 10 µm/s) are at least an order of magnitude higher than diffusion-limited polymerization rates of monomeric G-actin into F-actin ∼200 nm/s^92-94^, which drive filopodial and lamellipodial extension, suggesting that maximal cell migration speeds might require bleb expansion dynamics rather than F-actin self-assembly-driven protrusions to drive rapid leading edge advance.

The cell produces viscoelastic stresses on its surroundings as blebs expand/retract. The spatial structure of these stresses is examined in Figs. S4A (inset) and S4D, where high polymer stresses concentrate at the front of the cell, providing an additional resistance to pressure-driven bleb expansion. This results in lower cell displacements at the end of the bleb cycle for extracellular materials with a low stress relaxation time 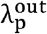Cell displacements at the end of the bleb cycle are greater in low density extracellular spaces (Fig. S3A) and are nearly insensitive to extracellular stress relaxation times (Fig. S3B). Low plasma membrane tension also causes maximal cell displacements in an individual bleb cycle (Fig. S3C), as membrane tension builds up at the cell front during bleb growth thus opposing further expansion (Fig. S4D) and a softer membrane provides less elastic resistance to forward plasma membrane motion. Overall, our model results show that larger cell displacements during a single bleb extension-retraction cycle are favored by low-density extracellular spaces, providing novel insights into the mechanisms used by phenotypically amoeboid cells when navigating complex porous networks. Likewise, our model demonstrates that soft actomyosin cortices, high myosin motor levels, and softer plasma membranes also favor cell displacement, identifying intracellular factors that may be engineered to design cells that move better in mechanically heterogeneous complex environments.

### Sustained adhesion-free bleb-based cell swimming in Newtonian environments requires cortical contractility oscillations

Since a single cycle of bleb extension-retraction results in minimal net cell displacement, we next explored whether bleb-producing cells can swim in adhesion-free conditions by producing bleb-based cell shape changes. To simulate more complex variations in the dynamics of cortex component amounts we introduce stochasticity in the kinetics of membrane-cortex linkers, actin and myosin, where membrane-cortex linkers stochastically associate in a force-independent manner and unbind by force with an effective dissociation rate that increases exponentially with force according to Bell’s law^95^ (see Supplemental Material for additional details). Hereinafter, we consider intracellular and extracellular spaces to be viscous fluids. We find that bleb-producing cells that maintain a consistently high contractility level cannot effectively migrate, as shown by the plateau reached by the sample average mean-squared displacement-time curve after a few bleb nucleation events (Fig. 3A and movie S2). The significant cell displacement achieved by the cells during the first bleb cycles suggests that cells could potentially migrate starting from a healed cortex with a fully pressurized intracellular cytoplasm. Therefore, we introduced a cortical contractility oscillator signal into the model, to mimic cortical contractility oscillations^96,97^, and allow the cortex to recover and “heal” before stochastically initiating the next bleb cycle. We find that cells are capable of modest swimming by producing high-to-low cortical contractility oscillations, where a high cortical tension phase characterized by multiple bleb nucleation events is followed by an intracellular pressure buildup recovery phase at low cortical contractility, resulting in mean-squared cell displacements that increase over time (Fig. 3B). Intermediate frequencies of the cortical oscillation signal seem to maximize cell swimming migratory capabilities (Fig. 3C). Overall, these results show that adhesion-free bleb-based cell shape changes combined with high-to-low cortical tensions are a possible strategy by which cells could move through biological tissues or fluids via a stochastic swimming behavior of repeated cycles of bleb-based protrusions driven by oscillations in cortical contractility. That is, our model predicts that amoeboid carcinoma and immune cell swimming is theoretically possible in Newtonian environments but is too slow to explain experimentally observed random motility coefficients associated with rapid T cell migration (∼10 µm /min^46,98,99^).

**Figure 3.**
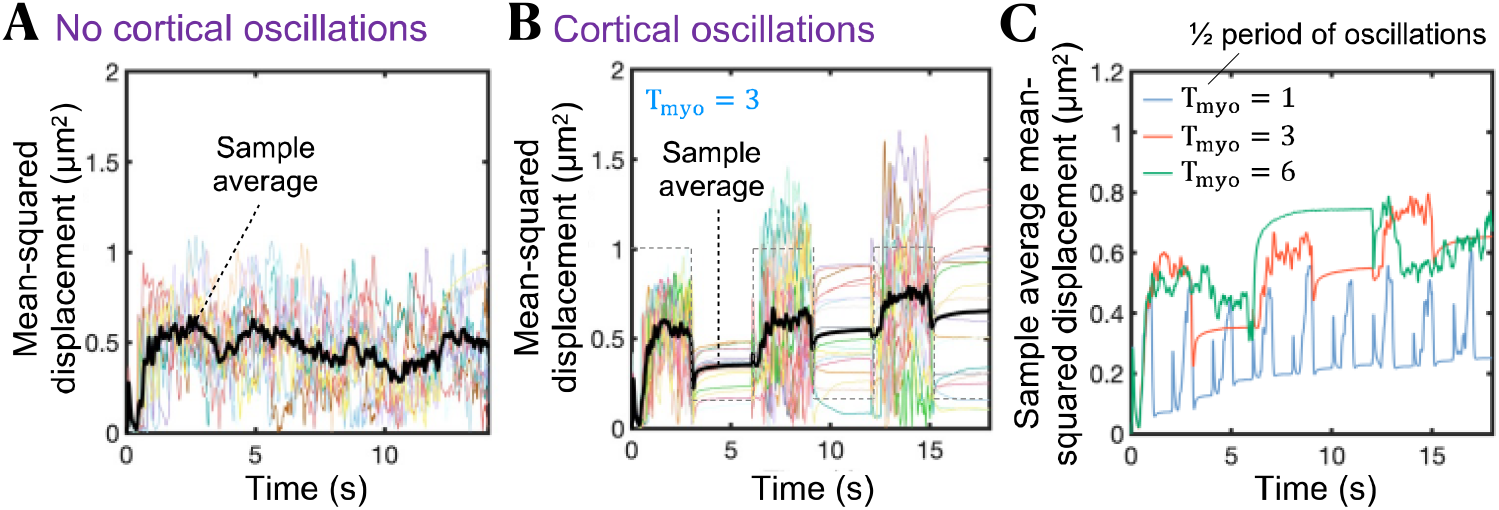
Cortical contractility oscillations are required for effective bleb-based cell swimming. (A,B) Time-evolution of mean squared displacement of 20 cells in the absence of cortical oscillations (A), and in the presence of cortical oscillations (B). The sample average over all cells (black line) increases over time for cells that produce cortical tension oscillations. Cells that do not produce cortical contractility oscillations are incapable of migrating effectively after the first bleb nucleation/expansion event. The gray dashed line in (B) indicates the waveform of the temporal oscillatory signal S _osc_. (C) Sample average mean squared displacement over time for three different periods (in seconds) of the cortical contractility oscillation signal. Intracellular and extracellular spaces are assumed to be Newtonian.

### Maximal bleb-based migratory potential achieved for high myosin motors and a soft cortex

To explore the optimum cellular conditions for maximal cell motility via the oscillatory bleb-based swimming mechanism identified above, we identify two important model parameters that strongly modulate cell motility: cortical stiffness and cortical tension. We find that a soft cortex is essential for cell motility (Figs. 4A and 4B). Cortical stiffnesses per unit of actin, κ_c_, higher than ∼0.2 pN/µm give rise to extremely low random cell motility coefficients, as shown in Fig. 4B. For an estimated number of actin units at each cortical node of 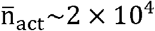 (see Supplemental Information), our model predicts that a cell with a cortical stiffness greater than 4000 pN/µm cannot swim effectively. Quantitative analysis shows a biphasic behavior of bleb nucleation frequency *ν*_bleb_ as a function of cortical stiffness. When cortical stiffness is high, the cortex is unable to rapidly adjust to deformations in the plasma membrane. This frequently places membrane-cortex linkers under high tension, leading to frequent membrane-cortex linker ruptures and increased bleb nucleation frequencies. In the limit of low cortical stiffness, in-plane cortical motion caused by contractility gradients can also often subject membrane-cortex linkers to a high tensional state causing frequent bleb nucleation events as well. Consequently, a minimum nucleation frequency is found at intermediate cortical stiffnesses, as depicted in Figs. 4C and 4D. Intermediate bleb nucleation frequencies correlate with maximal cell random motility coefficients (Fig. 4D). Whereas cells cannot swim under low-frequency conditions, since effectively cells nucleate a sequence of isolated bleb cycles, leading to no swimming, high-frequency blebbing cells are not capable of successfully pressurizing their cytoplasm between consecutive bleb events, causing a decrease in their swimming propulsion efficiency. Intermediate blebbing frequencies are therefore ideal conditions for swimming, where the synchronized nucleation of multiple blebs indeed allows cells to break time-reversibility and migrate with modest effectiveness (∼1 µm^2^/min) in the absence of adhesion-based forces. Interestingly, metastatic potential of prostate cancer cells have been reported to correlate with reduced F-actin and high blebbing frequency^100^. The polar angle between consecutive bleb nucleation events (see Supplemental Material for further details), here referred to as bleb nucleation correlation angle p_bleb_, monotonically decreases with cortex stiffness, as shown in Fig. 4E, with soft-cortex cells and stiff-cortex cells nucleating blebs at p_bleb_ ∼[60° − 70°] and p_bleb_ < 40, respectively, as depicted in Fig. 4F.

**Figure 4.**
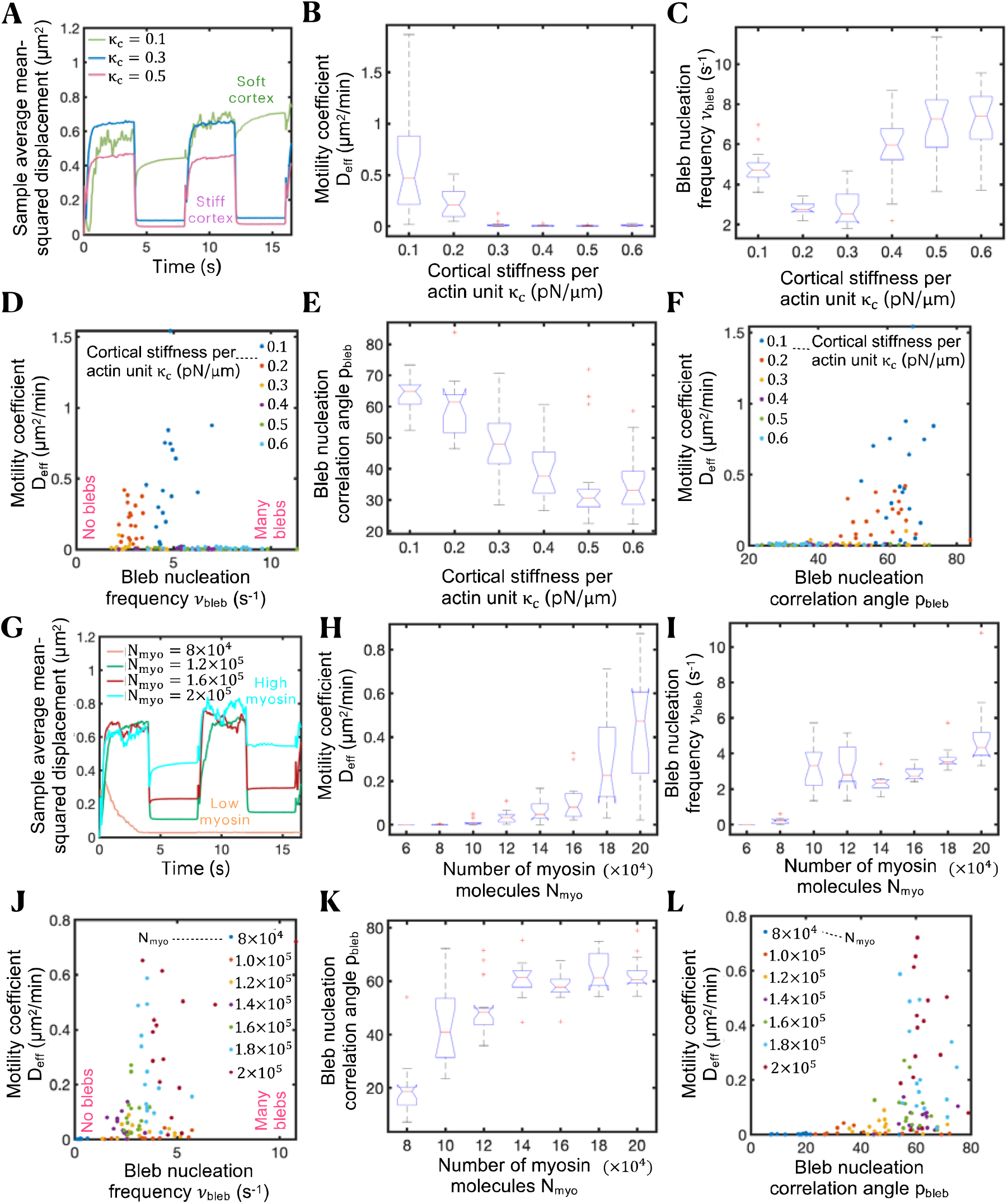
Swimming capabilities of bleb-producing cells can be modulated by cortical contractility and cortical stiffness. (A) Time-evolution of sample average mean squared displacement for four different values of the cortical stiffness per unit of actin parameter κ_c_ .(B,C) Effective random motility coefficient D_eff_ (B) and bleb nucleation frequency *ν*_bleb_ (C) as a function of κ_c_. Cells swim more efficiently for lower cortex stiffnesses. The effective diffusion coefficient D_eff_ is extracted from the data by fitting a line constrained to go through the origin in the mean-squared-displacement-time plots over 16 seconds. (D) Scatter plot showing the relationship between D_eff_ and *ν*_bleb_ for different values of κ_c_. Intermediate bleb nucleation frequencies correlate with maximal random motility coefficients. (E) Polar angle between consecutive bleb nucleation events p_bleb_ as a function of κ_c_. (F) Scatter plot showing the relationship between D_eff_ and p_bleb_ for different values of κ_c_. Maximal random motility coefficients are observed for polar angles of ∼60°-70°. (G) Time-evolution of sample average mean squared displacement for number of myosin motors N_myo_. (H,I) Effective random motility coefficient D_eff_ (H) and bleb nucleation frequency *ν*_bleb_ (I) as a function of N_myo_. Cells swim more efficiently for high number of motors. (J) Scatter plot showing the relationship between D_eff_ and *ν*_bleb_ for different values of N_myo_. (K) Polar angle between consecutive bleb nucleation events p_bleb_ as a function of N_myo_. (L) Scatter plot showing the relationship between D_eff_ and p_bleb_ for different values of N_myo_. Maximal random motility coefficients are observed for polar angles of ∼60°. Intracellular and extracellular spaces are assumed to be Newtonian.

We additionally find that high cortical contractility is essential for bleb-based cell motility, with high-myosin-motor cells possessing a higher motility coefficient than low-myosin-motor cells, as depicted by the mean-squared displacement curves, effective random motility coefficients and wind rose migratory cell track (Figs. 4G, H and S4, respectively). Higher number of myosin motors increases blebbing frequency and bleb nucleation correlation angle (Figs. 4I–L). Maximal cell motility coefficients not only correlate with intermediate blebbing frequencies (as already discussed) but also with intermediate bleb-to-bleb polar angles, as shown in the scatter plots in Figs. 4J and 4L. Maximal cell motility coefficients correlate with bleb nucleation frequencies of *ν*_bleb_ ∼4s^−1^ and bleb-to-bleb polar angles of p_bleb_ ∼60°. Cells that nucleate sequential blebs with low correlation angles (p_bleb_ < 40°), i.e. blebs on top of blebs, approach geometrical reciprocal motion resulting in vanishing cell motility coefficients, whereas cells that nucleate sequential blebs too separated from each other spatially (p_bleb_ > 70°) lose directionality/cell polarization, do not create effective non-reciprocal plasma membrane deformations and their migratory potential is penalized, as Fig. 4L shows. Whereas slow turnover time of cortex components, low fluid viscosity and low membrane tension favor bleb-based cell spreading, we have not observed any statistically significant difference in migration capabilities for the different values explored of cortex porosity, as depicted in Fig. S7. We hypothesize that the actomyosin cortical meshwork hinders cytoplasmic flows during both bleb expansion and bleb retraction, and so the slowing of cytoplasmic flows during both phases cancels out and its effect on bleb-based cell motility becomes insignificant. Overall, our model results show that cells with a soft cortex and elevated cortical tension exhibit more efficient motility by extending large bleb protrusions and the precise spatiotemporal nucleation of blebs, suggesting that engineering therapeutic T cells to possess increased cortical tension and/or a softer cortex would favor bleb-based T cell movement in complex environments. Notably, our cortical tension result is consistent with recent experimental findings, where high contractility pushed adhesive T cells into a faster amoeboid phenotype^21^.

### Blebs nucleate in regions of high cortical tension, low membrane-cortex linker density and high intracellular osmotic pressure

We next investigated the role of spatial variations in mechanical factors that potentially control the location of bleb nucleation sites, and thus bleb-based cell directionality. Bleb nucleation events are initiated by a local decrease in membrane-cortex cohesion. We explore three different scenarios: spatially dependent cortical myosin recruitment, spatially dependent membrane-cortex linker recruitment, and asymmetric distribution of electro-osmotic pumps within the plasma membrane. Blebs preferentially form in regions of high cortical contractility (high myosin) and low membrane-cortex linker levels, as shown in Fig. 5A, and in agreement with experimental observations^22,101,102^. Linkers preferentially reach a high tensional state in areas of myosin enrichment, as they are subjected to higher cortical inward forces that eventually cause them to dissociate by force in a runaway process, giving rise to local membrane-cortex detachment and bleb formation. As the bleb enlarges, the membrane-cortex detachment size keeps increasing due to the ongoing rupture of membrane-cortex linkers, while a new cortex forms beneath the plasma membrane. Concurrently, the expansion of the bleb shifts the cortical force at its neck from inward to outward, reducing the tensional state of linkers, and effectively preventing further detachment of the membrane from the cortex. In a similar way, linkers located in low linker density regions are overpowered by high cortical loads making them more susceptible to failure. The highly polarized orientational distribution of blebbing events forces cells to extend protrusions along one axis, restricting their motion to only one degree of freedom, and in turn impeding their migration as shown in Fig. 5B and consistent with the scallop theorem^88^. Therefore, enforcing the formation of blebs in a small subregion of the cell turns out to be a poor cell shape change-based swimming strategy. Linker failure is also exacerbated in high intracellular osmotic pressure regions (Fig. S8). The highly porous cortex does not sustain net osmotic forces, therefore an increase in local intracellular osmotic pressure causes membrane outward motion, straining linkers which results in a cascade of linker failures and bleb formation. This finding is consistent with the study by Yoshida & Soldati^103^, which showed that low osmolarity media triggers blebbing, highlighting the importance of osmotic stresses in blebbing. Overall, our model results show that a competition between membrane-cortex cohesion and a combination of osmotic and myosin-driven cortical forces can control the location of bleb formation sites, again identifying potential cell engineering design strategies to alter and control cell protrusion dynamics to enhance motility.

**Figure 5.**
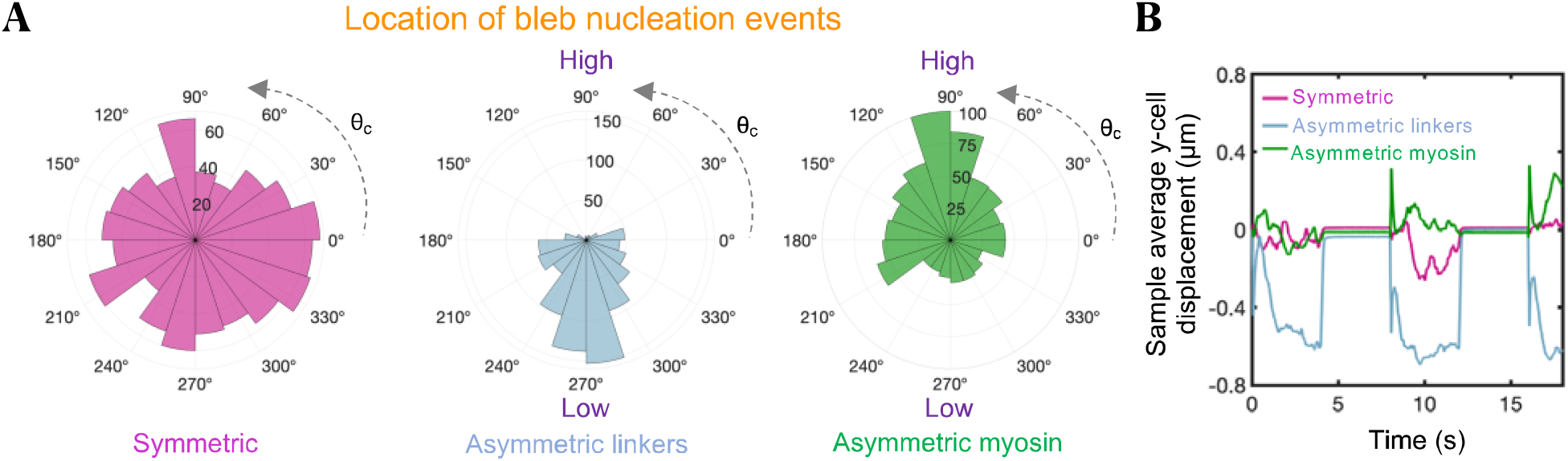
Blebs nucleate in regions of high myosin and low membrane-cortex linker levels. (A) Polar distribution of bleb nucleation events for three different conditions: uniform association of membrane-cortex linkers and myosin: 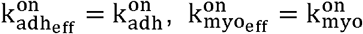 (red violet), asymmetric kinetic association rate constant of membrane-cortex linkers: 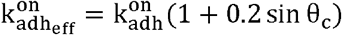 (blue) and asymmetric kinetic association rate constant of myosin: 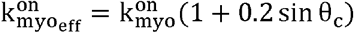 (green). Here, θ_c_ is the polar angle associated to each cortical node. (B) Time-evolution of sample average vertical cell displacements for the three different conditions. For the chosen parameter values, larger blebs are formed in the asymmetric linker configuration, resulting in larger displacements during the high cortical contractility phase. Intracellular and extracellular spaces are assumed to be Newtonian.

### Rapid cell migration requires cell-matrix adhesion forces and is potentially enabled by a hybrid bleb- and adhesion-based mode of migration for optimum cell motility

In order to analyze whether bleb-based swimming is a feasible mechanism that explains observed T cell motility *in vitro* and *in vivo*, we explored if blebbing alone can mediate rapid cell migration in the absence of adhesion-based forces, or if it requires adhesion interactions with the environment for optimal motility. We find that bleb-based coordinated cell-scale deformations are inefficient and lead to maximal migration motility coefficients of ∼1.5 µm^2^ /min (Fig. 4), an order of magnitude lower than reported *in vivo* cell motility coefficients of bleb-producing cells such as T cells (∼10 µm^2^ /min^21^), which display much faster speeds than those achieved by mesenchymal cells (∼1.5 µm^2^ /min for U251 glioma cells^104^), suggesting that swimming is not the primary T cell movement mechanism.

To address this discrepancy between maximum swimming speed and well documented T cell motility coefficients *in vitro* and *in vivo*, we investigated the effect of cell-environment adhesion interactions on the migration capabilities of bleb-producing adherent cells by constructing a simple hybrid adhesion-based, bleb-based cell migration biophysical model (Fig. 6A, see Supplemental Material for a detailed description of the model). Our findings indicate that adhesion interactions with the environment significantly increase cell migration speeds, with cells being capable of translocating approximately up to d_run_ ∼1.8 µm during an isolated bleb cycle of 9 seconds (Fig. 6B), for a mean speed of ∼12 µm/min. In this mode of migration, we observe that blebbing allows cells to push their cell front forward at a very fast rate (∼10 µm/s), much faster than F-actin polymerization rates and consistent with experimental observations^86^, followed by formation of focal adhesions at the cell front, which prevents cell rearward motion during bleb retraction and mediates subsequent traction force generation and fast forward cell translocation. Our model shows that bleb producing adherent cells migrate more effectively for slow cortex turnover times (Fig. 6B) and under high cortical tension conditions (Fig. S9A). Interestingly, while a softer cortex favors greater cell displacements during bleb expansion, a stiffer cortex causes larger cell displacements during bleb retraction, with the highest net cell displacements occurring at intermediate cortical stiffnesses (Fig. S9B). Repeated cycles of adhesion-based blebbing dynamics can lead to diffusive cell behavior, assuming that bleb nucleation sites are independent of the location of previous bleb nucleation events. We can again estimate the 3D effective cell motility coefficient associated with this hybrid migration mechanism as 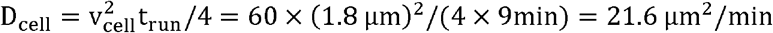. This value is comparable to the *in-vivo* motility coefficient of bleb-producing cells, suggesting that the hybrid cell migration mechanism is likely the mode of migration for T cells navigating through tissues.

**Figure 6.**
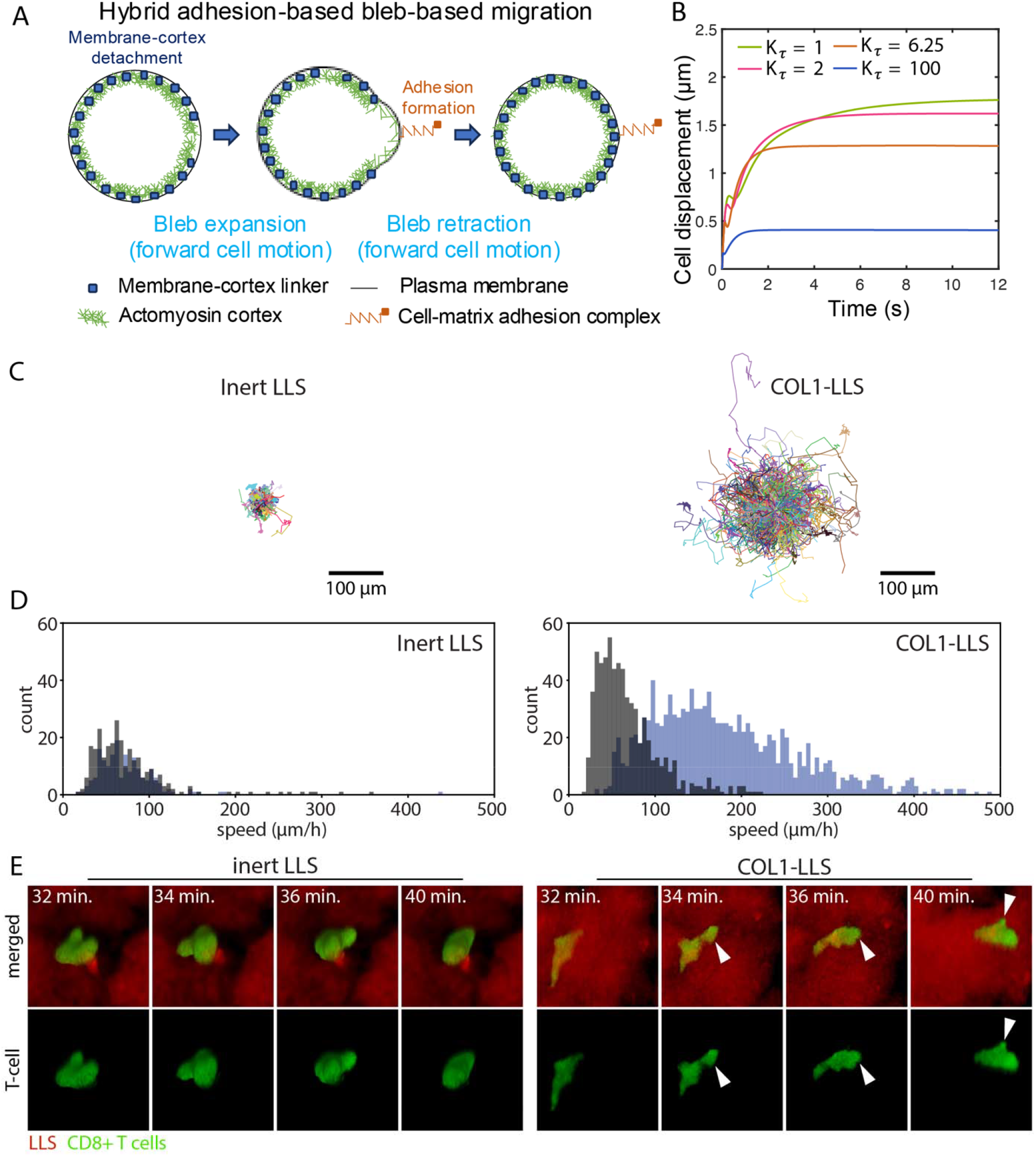
Biophysical model and experiments show that cell-extracellular matrix adhesion is required for rapid T cell motility. (A) Visual representation of one complete cell migration cycle in the hybrid adhesion-based bleb-based mode of migration. (B) Model prediction of the time-evolution of the cell centroid for four different values of the cortex turnover factor K_τ_. Larger cell displacements are achieved for slow cortex turnover times. Physiological cortex turnover times correspond to K_τ_∼1 (see Table S1). (C) Projected x-y migration trajectories of T cells in inert LLS microgels (left) and COL1-LLS microgels (right). T cells exhibit faster and more extensive migration in the presence of collagen. (D) Histogram of walking speeds for T cells migrating in inert and COL1-LLS. Speed was defined for each cell as the cell diameter divided by the time required to travel a distance equal to one cell diameter. Mean walking speed was calculated separately for two subpopulations: cells shown in blue reached a mean squared displacement (MSD) greater than one cell diameter squared within 3000 seconds of lag time, while cells shown in black did not meet this threshold. See Supplementary Movie S4 for experimental data corresponding to panels C and D. (E) Selected confocal time-lapse images showing a single T cell migrating through the 3D interstitial space between microgel particles in inert LLS (left) and COL1-LLS (right). See Supplementary Movies S5 and S6 for the full time-lapse sequences. White arrow indicates the bleb position.

To experimentally test our model prediction that cell-matrix adhesion forces are essential for rapid T cell motility, we employed established liquid-like solid (LLS) microgels with bioconjugated type 1 collagen (COL1)^105^ and examined human CD8^+^ T cell invasion in three-dimensional COL1-LLS environments^106^. Collagen conjugated to the surface of the microgels promotes cell adhesion and facilitates migration through the interstitial spaces between the microgel particles. This platform allows us to decouple MMP-dependent factors and specifically investigate the contribution of adhesion to 3D cell migration. The figure/supplementary video demonstrates that T cells are capable of squeezing through narrow interstitial spaces. However, this migration is significantly enhanced in the presence of adhesion cues on the microgel surface, compared to T-cell migration in inert LLS, where cells lack the ability to adhere to their surroundings. Taken together, our results demonstrate that T cell migration is significantly faster under adherent conditions in COL1-LLS compared to non-adherent conditions in inert LLS (Figs. 6C, D), supporting the presence of cell-environmental adhesion forces in promoting rapid motility.

Additionally, our model predicts that in cells employing a hybrid adhesion-bleb-based migration strategy, cell displacements remain unaffected by the stiffness of the substrate, as shown in Fig. S9C. This finding aligns with experimental observations from T cells on quasi-3D elastic platforms^21^. In these experiments, cells treated with Nocodazole, a microtubule destabilizer, exhibited high contractility and an amoeboid-like morphology. These cells showed a migration potential that was independent of the rigidity of the nanotexture (see Fig. 2C in reference^21^), potentially suggesting that Nocodazole-treated high contractile T cells might use a hybrid bleb-adhesion-based migration mode. The observed insensitivity of cell migration speed on substrate rigidity among these cells starkly differs from the behavior observed in the adhesion-based motor-clutch migration mode, where migration of highly contractile cells is sensitive to the stiffness of the substrate^25^. Crucially, the effectiveness of the hybrid adhesion-bleb migration mode hinges on the predominant formation of cell-matrix adhesions at the leading edge of the cell (Fig. S9D). If cell-matrix adhesions are largely formed at the rear of the cell, cell movement is predicted to be significantly impaired (Fig. S8D). Increased membrane tension at the leading edge (Fig. S4D) may locally activate adhesion molecules^107^, such as integrins, thereby promoting a biased formation of cell-matrix adhesions at the front. This spatial bias would enhance directed migration by coordinating hydrostatic pressure gradients with traction forces. We also note that enhanced traction forces mediated by actin polymerization forces have not been considered in this analysis, but it is predicted that they would enhance significantly the cell migratory potential of adherent blebby cells. However, how cells coordinate the spatiotemporal assembly and disassembly of focal adhesions, actin polymerization and traction forces with blebbing remains to be determined.

In summary, our findings uncover potential mechanisms governing the remarkable ability of T cells to rapidly navigate through tissues, suggesting that cell-matrix adhesion enables these cells to generate rapid and sustained motility within complex microenvironments. Our model and experimental results do not support an adhesion-free swimming strategy relying solely on changes in cell shape, specifically through the formation of protruding blebs. Instead, this mechanism appears to allow them to slowly explore/sample the surrounding microenvironments prior to robust directed movement. Importantly, our findings suggest that fast migration requires the presence of cell-surrounding matrix adhesion interactions. The synergy between fast forward plasma membrane motion driven by bleb expansion, followed by cell-matrix adhesion formation and subsequent traction force production characteristic of mesenchymal migration, likely optimizes T cell motility.

## DISCUSSION

We developed a mechanistic model of bleb-based cell motility and conducted T cell migration experiments in soft, liquid-like solid granular media under both adherent and non-adherent conditions. Our work elucidates the potential physical principles and molecular components that modulate T cell migration, examines whether adhesion is required for rapid motility, and explores how perturbing these molecular regulators can modulate their migration potential. The amoeboid cellular phenotype has been associated with immune cell migration^21,22,108^ as well as with multiple cancer cells displaying epithelial to amoeboid transitions with enhanced cell invasion, immunosuppression, stem cell potential and metastatic capabilities^109-113^. Although the developed model is aimed at providing insights to effectively engineer therapeutic T cells with enhanced migration capabilities, the model’s applicability extends broadly to all bleb-producing cells, including amoeboid cancer cells, with a major goal being to induce fast amoeboid T cell migration and target/inhibit amoeboid cancer cell migration. While our model addresses crucial aspects relevant to this research area by elucidating the most plausible mechanisms of fast T cell migration, consistent with experimental data, and within the constraints of current modeling capabilities and knowledge, it does not capture every complex detail inherent in real cells. Integrating every molecular constituent and biological aspect into our biophysical model is beyond practical reach. Missing explicit intracellular and extracellular factors include microtubules, septins, aquaporins and cytokines, which are known to influence cortex behavior, the functionality of cell adhesion molecules, and the overall migration of T cells^21,114-116^. According to Tabdanov et al.^21^, microtubules play a more significant role in regulating GTPase RhoA signaling than serving as direct mechanical elements. GEF-H1 associates with polymerized microtubules, and upon depolymerization, GEF-H1 is released, increasing its guanosine nucleotide exchange activity. This leads to higher RhoA activation and increased cortical contractility^21,96^. Our model indirectly incorporates the effect of microtubules on T cell migration by simulating this global/local effect on actomyosin contraction (Figs. 2D and S9A). Additionally, our model does not account for the role of the nucleus, which acts as a physical barrier and completely arrests T cell migration through interstitial pores smaller than 7 µm^2^ when matrix metalloproteinase-independent pathways are involved^117^. While the nucleus is thought to impede migration by resisting cytoplasmic flows, due to both steric hindrance and mechanical resistance through its binding to the cytoskeleton, deformation of the nucleus under confinement may also activate mechanotransduction pathways that enhance actomyosin contractility^118,119^. Although such effects could influence migration speed, explicitly modeling the nucleus would require assumptions about its material properties and coupling to the cortex that remain speculative. We therefore chose to focus on well-characterized mechanical features while acknowledging the nucleus as an important future direction. Overall, we expect that mechanically the nucleus will act to slow down migration, and therefore our analyses omitting the nucleus can still be accurately regarded as theoretical upper limits.

Another important factor not explicitly captured is the effect of geometric confinement and wall-like boundaries. In vivo, T cells frequently encounter structures such as matrix pores or surrounding cells, which may impose physical constraints that alter migration mechanics. Confinement may facilitate directional bleb expansion along narrow channels but can also hinder migration during bleb retraction through nonspecific frictional interactions, particularly involving the larger surface area of the cell body. This is consistent with our finding in Fig. S9D, which shows that the hybrid adhesion-bleb migration mode is most effective when cell-matrix adhesions are concentrated at the leading edge. Under confinement, this spatial distribution may be disrupted, potentially slowing net migration. Simulating these interactions would require incorporating boundary forces at high spatial and temporal resolution while conserving momentum, a challenge that remains computationally infeasible in our current framework. As discussed in our recent review^120^, the physical nature of these weakly-adhesive interactions is still poorly defined and may involve weak clutches mediated by non-specific van der Waals forces, hydrogen bonds, ionic interactions, or components of the glycocalyx. Future modeling and experimental efforts will be needed to fully understand how confinement shapes T cell motility. In addition, our model can be extended in the future to incorporate more realistic features such as aquaporin dynamics, local changes in actin cortex permeability, and electroosmotic effects. These additions, however, were beyond the scope of the present study.

Our model results indicate that sustained adhesion-free bleb-based cell motility requires oscillatory cortical forces in Newtonian environments, where cells alternate between a high-contractility motility phase and a low-contractility pressure buildup phase. These pulsatile actomyosin forces, which can arise from different mechanochemical sources, including calcium signal fluctuations^97,121^, myosin phosphorylation-based biochemical oscillators^122,123^, and ERK-based mechanochemical signaling, among others, do not seem to be essential in bleb-based cell swimming only, but also in other biological contexts including adhesion-based cell migration^96^, compaction^124^, cell intercalations^97^ and tissue integrity^123^ during morphogenesis, and temporal regulation of actin polymerization and vesicle exocytosis^121^. Future theoretical work could incorporate existing reaction-diffusion models^125-127^ into our bleb-based cell migration framework to more explicitly capture the biochemical mechanisms underlying contractility oscillations. For example, integrating models of oscillatory calcium dynamics could help elucidate how biochemical calcium signaling networks contribute to the temporal regulation of cortical tension and its impact on cell motility.

Repeated sequences of a single bleb nucleation event, bleb expansion, cortex recruitment and bleb retraction do not cause sustained cell swimming in Newtonian fluids, since the plasma membrane follows similar paths during bleb expansion and retraction. However, we show that individual blebs in viscoelastic environments or spatiotemporal synchronization of multiple bleb nucleation events in Newtonian environments allows cells to break time-reversibility and effectively migrate in the absence of adhesion-based forces (i.e. swim), albeit at rates well below those exhibited experimentally by T cells. Additionally, our results indicate that coordinated cell-scale deformations driven by blebs are inefficient and result in maximal motility coefficients of ∼1.5 µm^2^ /min, an order of magnitude lower than those reported by bleb-producing cells *in vivo* (∼10 µm^2^ /min^21^). In such poor swimming conditions, interstitial flows would dominate in certain tissues over the inherent swimming motion of cells rendering cell swimming less relevant to cell translocation^128,129^. Our analysis gives a new physical perspective of the mechanism underlying potential swimming of bleb-producing cells and provides a strategy to modulate cell swimming potential by targeting the cortex, plasma membrane and extracellular environment.

Other possible adhesion-free cell swimming mechanisms rely on peristaltic waves^130^, osmotic pressure gradients^131^, Marangoni stresses^49,50^ or slippage between plasma membrane and surrounding fluid^132^ possibly caused by mechanical coupling of transmembrane proteins, powered by F-actin retrograde flows, with the surrounding fluid^40,48^. Although these migration models are not analyzed within our theoretical framework, our experimental results already indicate that adhesion-based migration is inherently faster and more efficient than non-adhesive modes. Reversat et al.^44^ proposed an alternative mechanism for adhesion-free migration, suggesting that T cells can navigate through serrated microfluidic channels without relying on adhesion or friction-based forces. Their theory posits that cell migration is driven by intracellular pressure gradients generated by retrograde flows and the curvature of actin flows caused by the channel wall topography. While their experimental results show an intriguing dependence of cell migration speed on wall topography, their proposed migration model contradicts established physical laws. The authors claim that the local force exerted by intracellular hydrostatic pressure gets directly transmitted to the channel walls. According to their theory, this pressure force, acting perpendicular to the walls, and summed across the cell’s surface, generates a net hydrostatic force, serving as a propulsion mechanism that drives the cell in the direction opposite to the actin flows. Their interpretation overlooks critical physical aspects of plasma membranes. A local increase in intracellular hydrostatic pressure will induce a small efflux of water, drawing the plasma membrane inwards, away from the channel walls, thus not transmitting any hydrostatic force to the walls, as the authors claim. In addition, it is unclear whether T cells migrated through these microchannels in the complete absence of adhesion/friction forces, as the authors claim, due to the technical challenge of completely blocking nonspecific adhesions. A plausible interpretation of their experimental findings, consistent with our model results, is that the magnitude of the adhesion/friction-based forces with the cellular environment (i.e. channel walls) may depend on its topographical features. We note that weak adhesions that last only a few seconds are sufficient in our model to enable fast migration in the hybrid amoeboid-mesenchymal mode.

Although cell signaling might play an important role in the coordination of multiple blebbing events, cells can orchestrate blebbing and control bleb frequency and bleb-to-bleb distance by temporal modulation of cortical tension and stiffness, where high tension and soft cortex favor optimum conditions. To evaluate whether higher speeds could be achieved under non-adhesive conditions, we explored non-physiological parameter combinations, such as extremely soft cortices paired with high myosin levels. While these conditions transiently increased speed, they rapidly caused mechanical instabilities, including cortical rupture and loss of shape integrity, making them unsuitable for sustained motility. Notably, the observed increase in speed with higher cortical tension aligns with experimental findings that associate elevated contractility with rapid amoeboid-like migration^5,24,133^. Motor activity levels can be modulated by the internal cellular state as well as by environmental cellular conditions. As an example, nuclear deformation mediated by enhanced cellular confinement can activate mechanotransduction pathways^118,119^ that enhance actomyosin contractility and therefore would be predicted to increase cell swimming potential. Whereas low contractile cells nucleate blebs at low frequencies and in regions near recent bleb nucleation sites, high contractile cells nucleate blebs at ∼4Hz and cause membrane-cortex detachment at bleb-to-bleb separation polar angles of ∼60. These blebbing frequencies and bleb-to-bleb separation distances correlate with maximal cell motility coefficients.

In addition, our model shows that blebbing events are expected to be favored in cell regions of high cortical forces, low membrane-cortex adhesion and local intracellular osmotic pressure. This is consistent with experimental data, where blebbing was less frequent in the ezrin/radixin/moesin-rich uropod-like structure of cells^28,134-136^ and favored in calcium-dependent high myosin contraction regions at the leading edge of cells^13^. Chemoattractants, which have also been associated with bleb polarization^137-139^, potentially modulate some of the three driving factors mentioned above. Interestingly, we find that unidirectional bleb-based cell swimming is not possible via highly polarized bleb nucleation events, strongly suggesting that persistent directional cell migration, observed in different confined systems^11,26,43,140^ might require the presence of cellular frictional/adhesive forces with the environment, as assumed in the hybrid amoeboid-mesenchymal model^39,44^.

Our findings indicate that T cells may use a hybrid migration strategy, integrating bleb formation with cell-matrix adhesion to optimize motility. Our model is supported by experimental T cell migration data, which demonstrate that T cells move significantly faster in microgels functionalized with type 1 collagen than in inert gels. In our system, collagen monomers are present only on the surface of the microgels at the cell-gel interface. The 3D platform is composed of polyacrylamide microgel ensembles, where the interstitial spaces between particles provide physical pathways for cell migration. This approach uniquely enables independent control over structural parameters, including particle size, water content, stiffness, topography, topology, and permeability, while enabling separate modulation of COL1-mediated adhesion, as previously described^141,142^. For this study, we employed two conditions: collagen-conjugated LLS (COL1-LLS) and inert, unconjugated LLS microgels. The inert gels, lacking adhesion molecules, have been shown to create frictionless environments^141^, offering the most direct and effective negative control for adhesion-based migration. We have demonstrated that fully hydrated polyacrylamide exhibits a negligible friction coefficient^141-143^, and therefore does not provide adhesive cues to migrating cells. In contrast, COL1 bioconjugation introduces adhesion sites that support cell binding and promote migration along interstitial paths^105,106^. Accordingly, this experimental design provides a robust framework to distinguish between adhesion-dependent and adhesion-independent T-cell migration.

Unlike in bleb-based cell swimming, the cell advances in the same direction during both bleb expansion and retraction phases in this proposed hybrid mode of migration, achieving speeds that are comparable to physiological T cell migration speeds. Our results indicate that adherent blebby T cells can exhibit a highly migratory phenotype by increasing their contractility and fine-tuning their cortical stiffness to intermediate values. This mode of migration is consistent with a recent study that shows that T cells push and pull on the extracellular matrix and require adhesion-based forces to migrate through *in vitro* systems^52^. Recent experimental observations additionally support this hybrid mode of migration. For instance, melanoma cells exhibit blebbing while moving through soft collagen matrices, pushing collagen aside at the leading edge during bleb expansion, and pulling it in during bleb retraction via adhesions^144^, as in our hybrid model. Similarly, rapid β1 integrin-dependent bleb-based migration of breast cancer cells involved extracellular matrix reorganization at membrane blebs^145^ with integrin clustering occurring at these sites. These findings align with the fast hybrid adhesion-based bleb-based migration mechanism, which relies on the predominant formation of cell-matrix adhesions at the cell’s leading edge. It is also consistent with another cell migration study, where recruitment of the branched actin nucleator Arp2/3 frequently follows membrane-cortex detachment, depletion of the ERM membrane-cortex linker ezrin and protrusion expansion^102^. In some instances, however, cell-extracellular matrix adhesion formation and blebbing events anticorrelate in space, potentially suggesting the presence of F-actin assembly-driven protrusions or pressure-only-driven protrusions^146^. Interestingly, recent work by Te Boekhorst et al.^147^ demonstrated that cancer cells can exhibit mixed migration modes combining blebbing and actin-based protrusions, with amoeboid phenotypes prevalent under low-adhesion conditions, consistent with effective migration using a hybrid mesenchymal-bleb-based mechanism.

Recent modeling results also suggest that cortical squeezing at the cell rear combined with focal adhesion formation at the cell front is a fast and robust cell migration strategy^148^. It is unclear though how blebbing, actin polymerization forces, traction forces, and spatial dynamics of focal adhesion assembly and disassembly integrate all together in this model.

Experimental studies in mesenchymal cells that use the motor-clutch model showed that clutch-extracellular matrix binding and initial force transmission can occur within 1–10 seconds^149,150^. This timescale is comparable to the duration of a typical bleb cycle (Fig. 6B, green curve). In the context of bleb-driven migration, actin cortex reassembly begins rapidly following bleb expansion, and myosin II recruitment typically occurs within 5–10 seconds^151,152^. Therefore, effective adhesion-mediated migration during bleb retraction does not require adhesion formation to be instantaneous after actin recruitment, but rather only needs to occur within seconds following cortex assembly, approximately within the same timescale as myosin recruitment. In our hybrid bleb-based mesenchymal model, cell-matrix adhesion is assumed to form at the end of bleb expansion, a timing that is consistent with this sequence of events. This assumption of immediate adhesion and traction after expansion defines an upper bound on cell migration speed in our model. Future extensions of our model will aim to explicitly incorporate actin polymerization-driven protrusions and dynamic adhesion kinetics to more fully characterize the parameter space and transitions between swimming, bleb-based adhesion-dependent migration and mesenchymal-like migration.

Although friction has been associated to the major factor explaining cell migration in passivated microfluidic devices^11,140^, it remains uncertain whether cellular interactions with the extracellular matrix in healthy or transformed tissues exhibit more elastic-like or friction-like characteristics over long time scales. Notice that, while our model assumes that the mechanical interactions with the extracellular environment are elastic in nature, it can, in effect, account for friction-like interactions with the environment. This is supported by the fact that cell-matrix elastic interactions are effectively equivalent to friction-like cell-matrix interactions under conditions of high dissociation rates of cell-matrix adhesion bonds^153,154^. Therefore, this bleb-based friction-based migration regime can be achieved by cells embedded in extracellular matrix within in vitro or in vivo environments and falls under the bleb-adhesion-based hybrid framework we outline here.

Overall, our work provides a new physical perspective of the possible mechanisms underlying the migration of blebby cells such as T cells and some cancer cells. The developed model provides not only strategies to enhance exploratory bleb-based T cell swimming but also suggested that pushing cells into a hybrid bleb- and adhesion-based mode of migration is a promising engineering strategy to increase T cell migration in tumors. Moreover, our findings suggest that antifibrotic therapies, targeting fibrotic tumor environments, can have contrasting effects on T cell migration, either slowing or enhancing movement depending on the specific treatment, emphasizing the importance of controlled therapeutic interventions. While dense, small-pore matrices may hinder T cell infiltration, intermediate matrix densities can provide the necessary adhesion sites for T cells to grip onto the environment and migrate more effectively.

Additional studies are needed to investigate to what extent this hybrid mode of migration can be adopted by T cells *in vivo*, which presents an exciting avenue for prospective theoretical and experimental research. To achieve this, advanced imaging, cell engineering and quantitative analysis combined with biophysical modeling will be indispensable in uncovering cell-specific and context-specific migration mechanisms. Ultimately, a faithful model can be used to define optimal states for maximal T cell migration and maximally effective immune cell therapies, and thereby guide the development of genetically engineered T cells optimized to migrate through mechanically challenging tumor microenvironments.

## METHODS

### Preparation of Collagen-Coated and Inert Poly(acrylamide-co-acrylic acid) Microgels

The bioconjugated polyacrylamide microgels were synthesized as previously described^105,106^. In brief, collagen-conjugated and inert (unconjugated) poly(acrylamide-co-acrylic acid) microgels were fabricated through free radical polymerization using acrylamide monomer, N,N′-methylenebisacrylamide crosslinker (bis-acrylamide, Sigma-Aldrich, Cat# 1003520262, St. Louis, MO, USA), pure acrylic acid (Thermo Scientific, Cat# L04280.36, Waltham, MA, USA), and Acryloxyethyl thiocarbamoyl rhodamine B (Sigma-Aldrich, Cat# 908665, St. Louis, MO, USA). Ammonium persulfate (APS, Sigma-Aldrich, Cat# 248614-100, St. Louis, MO, USA) and TEMED (Fisher Scientific, Cat# B9150-100, Waltham, MA, USA) were added as initiators. The polymerized hydrogel was then mechanically ruptured to create microgel particles. Carboxylic acid groups were activated with EDC (Gibco, Cat# BC25-25, Thermo Fisher Scientific, Waltham, MA, USA) and NHS (Alfa Aesar, Cat# A10312.22, Haverhill, MA, USA) in MES buffer (pH 5.5, Fisher Scientific, Cat# BP300-100, Waltham, MA, USA). Activated microparticles were conjugated with type I bovine collagen (Advanced Biomatrix, Nutragen, Cat# 5010-50, San Diego, CA, USA) at neutral pH. Unreacted sites were quenched with ethanolamine (Sigma-Aldrich, Cat# E6133, St. Louis, MO, USA). Microparticles were thoroughly washed with PBS (1X, Corning, Cat# 21-040-CV, Corning, NY, USA) before use.

### T Cell Maintenance

Purified human CD8^+^ T cells (iQ Biosciences, Alameda, CA) were grown in RPMI-1640 supplemented with 10% fetal bovine serum and 2 mM L-glutamine. Recombinant human IL-2 (BioLegend. Cat#589104, San Diego, CA) at the concentration of 100 ng/mL was added to the media for the maintenance of T cells. For activation, T cells were seeded at 3 x 10^6^ cells/mL in 6-well cell-culture treated plates in media containing soluble human anti-CD3 (1 µg/mL) and anti-CD28 (5 µg/mL) antibodies (BioLegend, San Diego, CA) for 48 h.

#### Long-term microscopy assay

Prior to the experiment, T cells were harvested and labeled with CellTracker™ Green CMFDA according to the manufacturer’s instructions. An equal number of labeled cells were suspended in 2 groups, collagen-coated and inert microgels, and then seeded into a 96-well plate that has a type #0 coverslip for optimal imaging. Imaging time-lapses were acquired as z-stacks (4um step size at 1024x1024) at 3-minute intervals using a Nikon AXR confocal microscope (Nikon Instruments, Inc USA). The ultra-high sensitive Nikon Spatial Array detector (NSPARC) and high-speed resonant scanning mode were also employed to optimize spatial and temporal resolutions while maintaining low cellular phototoxicity. Cells were maintained at 37°C and 5% CO_2_ in a stage-top incubation chamber (Aurita Biosciences), and timelapse images were acquired with a 10X Plan APO Lambda D air objective (NA 0.45).

#### Cell Segmentation, Tracking, and Analysis

A maximum intensity projection was created from z-stacks for each sample. The inherent non-isotropic motion/drift was compensated by optical flow to ensure that the detected motion was referenced to the LLS microgels. The cells were segmented with the Cellpose algorithm^155^. The centroids of single-segmented cells were tracked with an Image J Linear Assignment Problem (LAP) tracker algorithm. The individual mean squared displacements (iMSD) and mean speeds for each track were calculated, with short detections and those below-the-noise floor filtered out. The histogram of the mean speed of the tracks was plotted, labelling the two distinct populations revealed by the FPT histogram.

## ACKNOWLEDGEMENTS

Research reported in this publication was supported by grants from the National Institutes of Health U54CA210190, P01CA254849 and U54CA268069, from the National Science Foundation, grant number 2222434, and partially supported by the Nikon Center of Excellence at the H. Lee Moffitt Cancer Center. We thank members of the Provenzano and Odde labs for helpful conversations throughout the course of this work. The content of this work is solely the responsibility of the authors and does not necessarily represent the official views of the NIH or NSF.

---

**Figure 1A.**
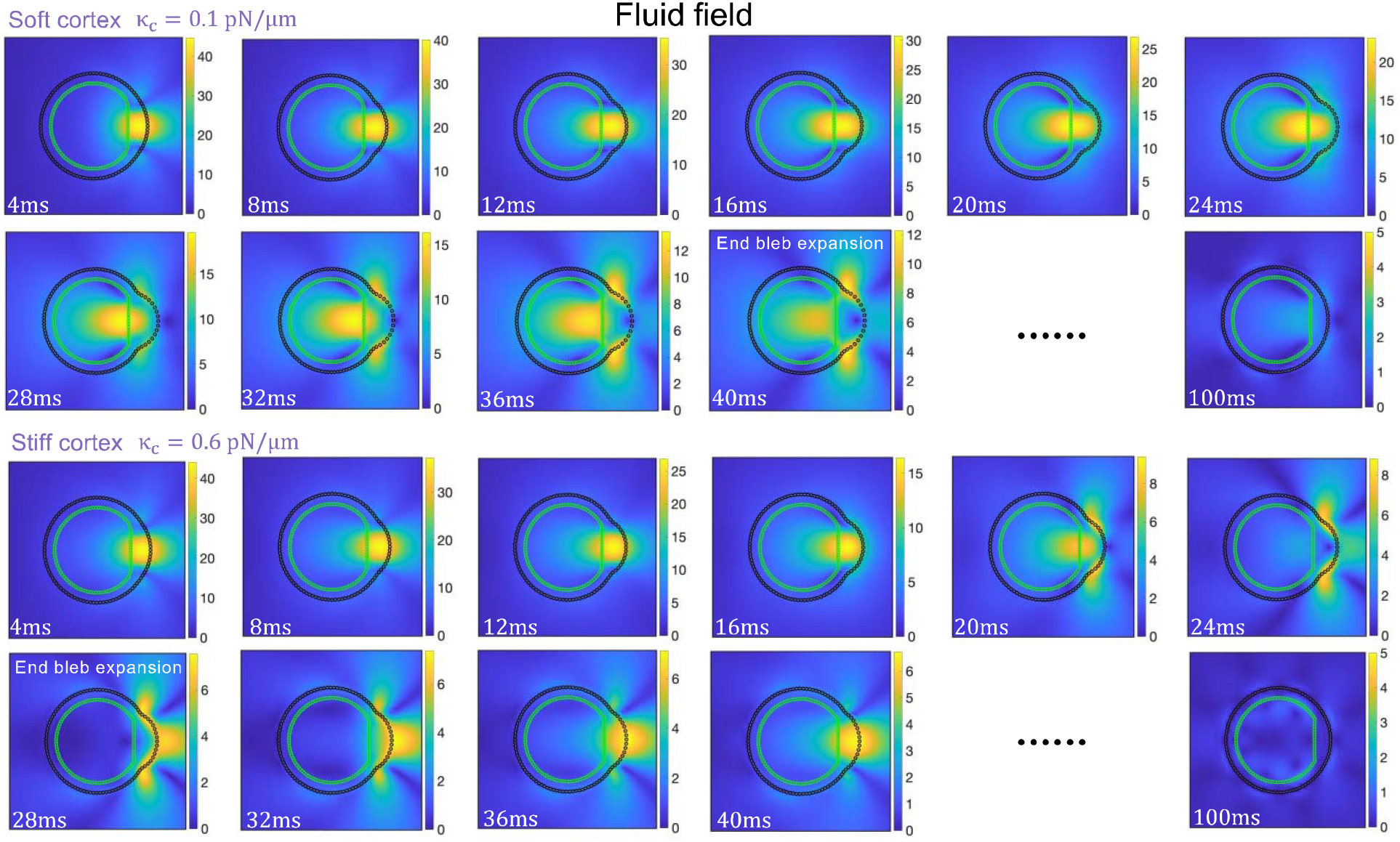
Magnitude of the fluid field (μm/s) throughout a bleb cycle for two different values of the cortex stiffness per unit of actin parameter κ_c_. Higher fluid velocities are observed for a softer cortex during both bleb expansion and bleb retraction. Membrane-cortex linkers are broken by hand locally in a small region at time t = 0, and are not allowed to break thereafter.

**Figure 1B.**
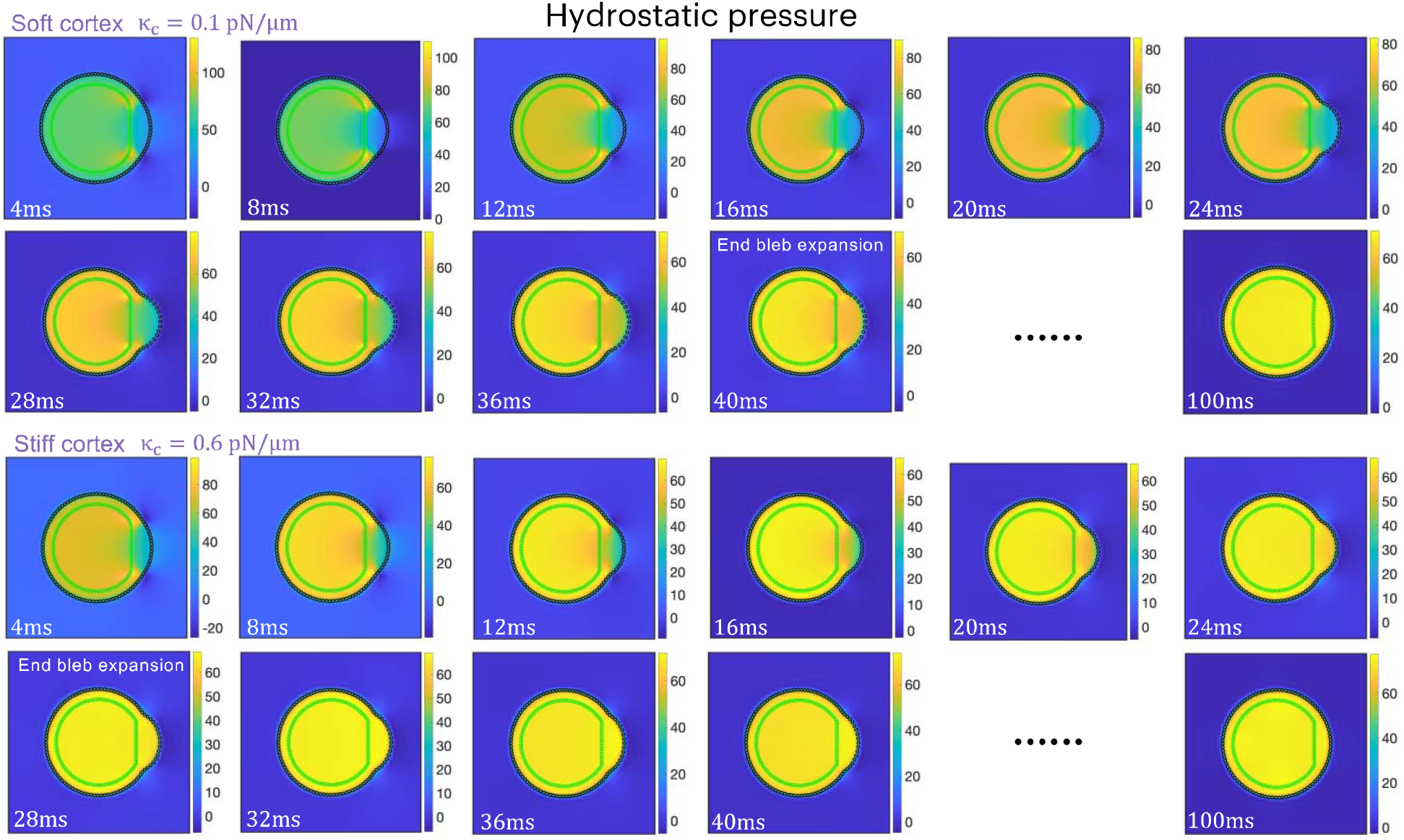
Magnitude of the hydrostatic pressure (pN/μ m) throughout a bleb cycle for two different values of the cortex stiffness per unit of actin parameter κ_c_. Hydrostatic pressure equilibration and the end of bleb expansion occur at earlier times in the bleb cycle for a stiff cortex. Membrane-cortex linkers are broken by hand locally in a small region at time t = 0.

**Figure S2.**
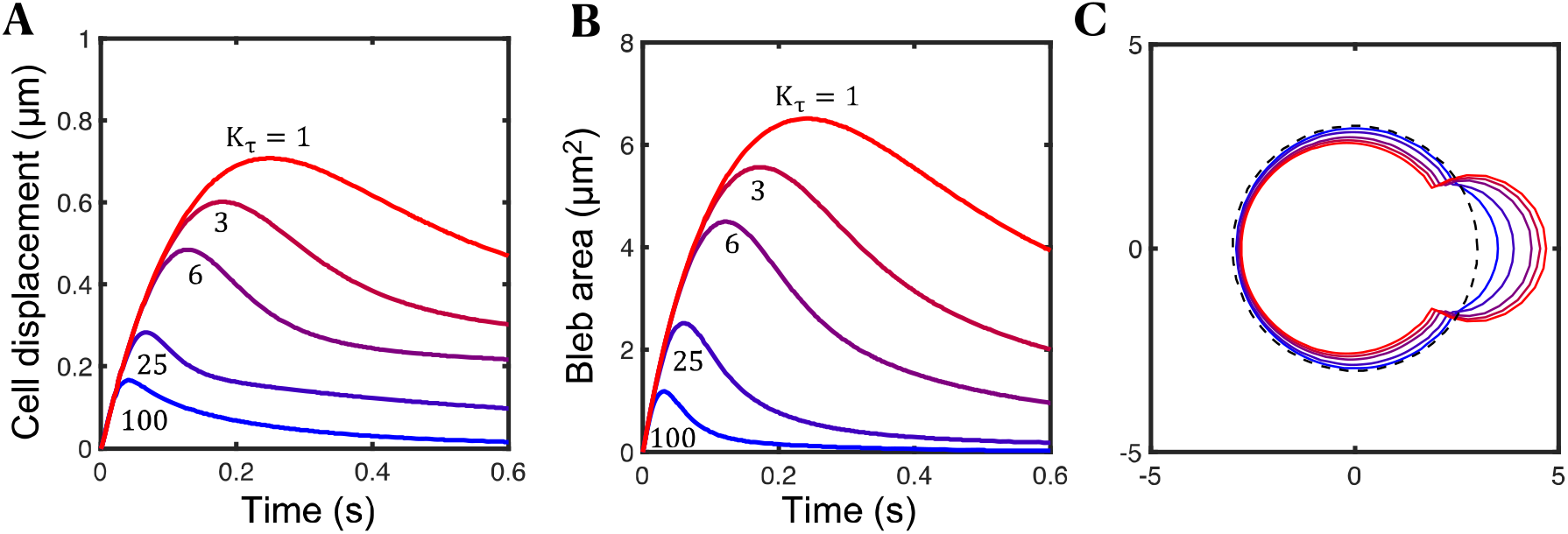
Cortex turnover regulates bleb retraction dynamics. (A–C) Time-evolution of the cell centroid (A), time-evolution of the bleb area (B) and cell shape at maximum bleb area (C) for five different values of the cortex turnover factor K_τ_. Extracellular and intracellular spaces are considered Newtonian fluids.

**Figure S3.**
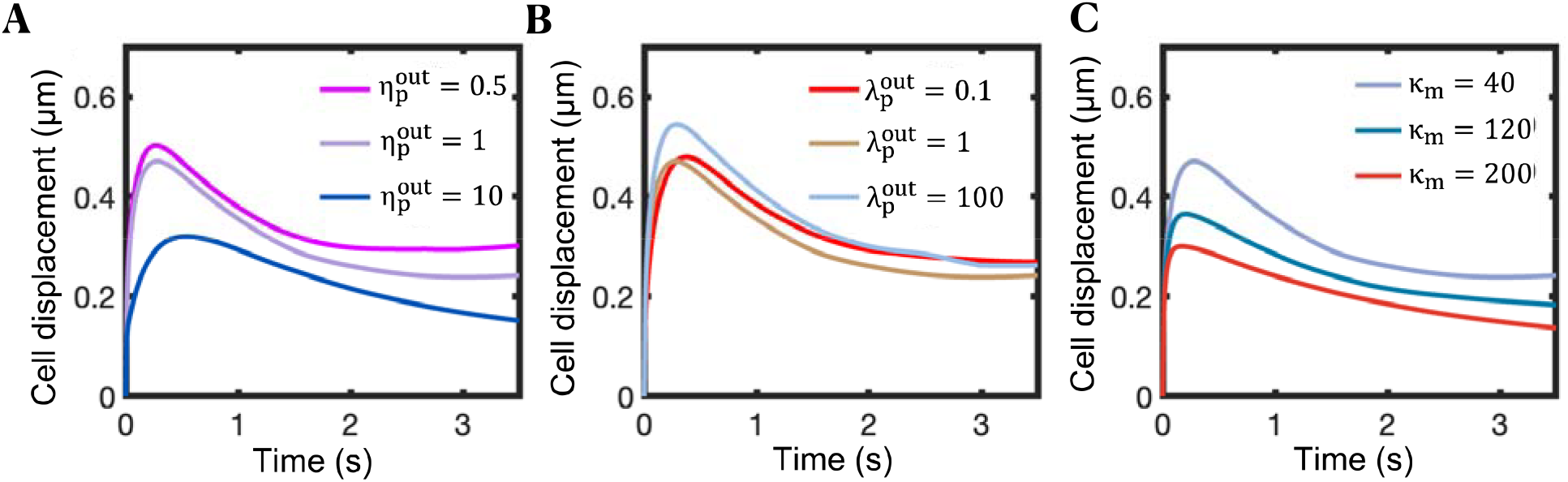
Cell displacements during a bleb cycle are influenced by extracellular and plasma membrane mechanical properties. (A–C) Time-evolution of the cell centroid for three different values of the cortex turnover parameter K_τ_ (A), three different values of the extracellular stress relaxation time 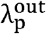 (in seconds) (B), and three different values of the of the plasma membrane spring stiffness 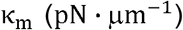 (C). In all panels, membrane-cortex linkers are broken by hand locally in a small region at time t = 0, mimicking a local downregulation of membrane-cortex linkers due to a chemical signal or force. Unless otherwise specified, polymeric model parameter values: 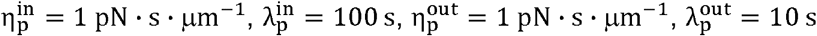.

**Figure S4.**
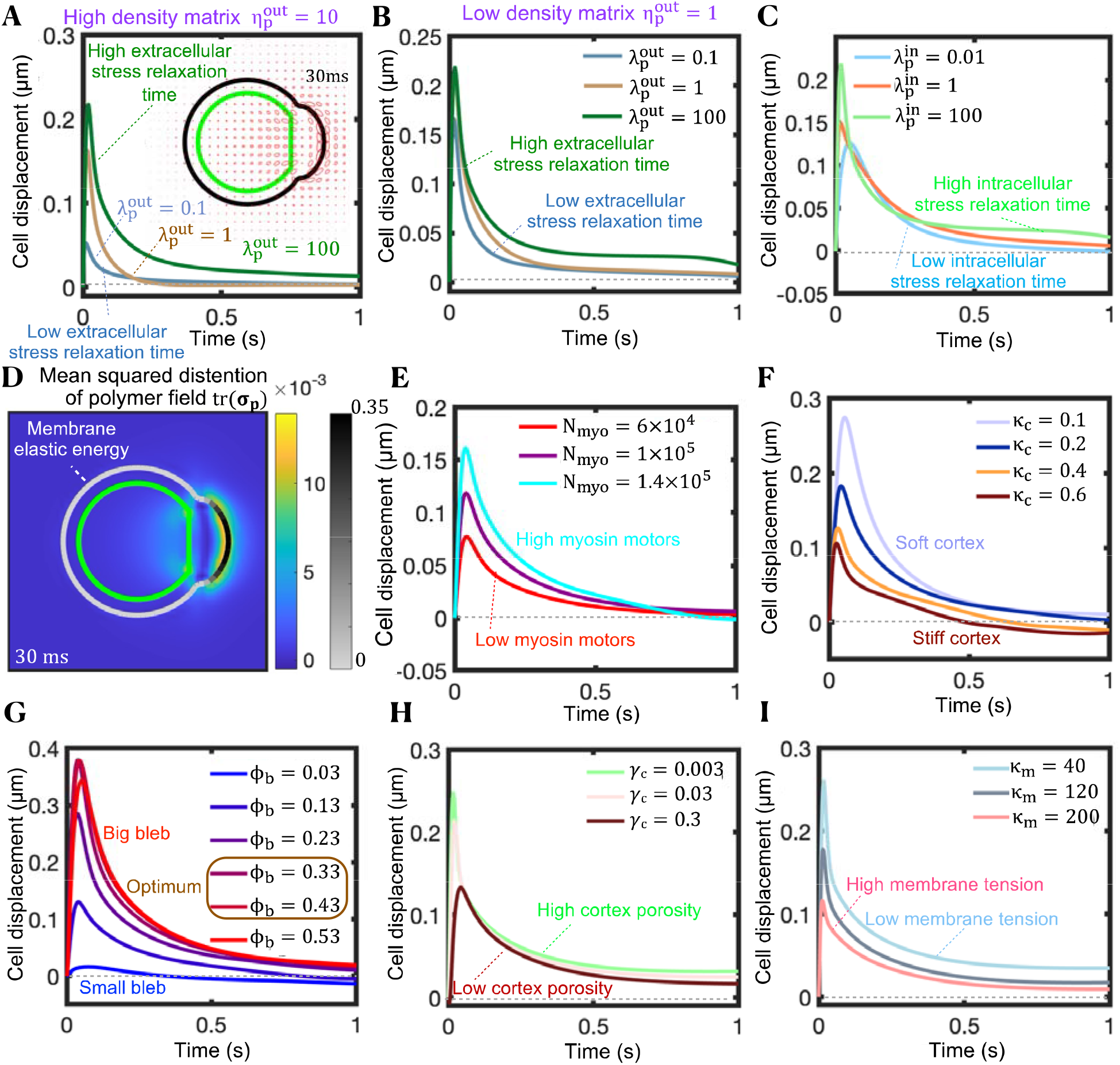
Net cell displacements for fast turnover kinetics during a single bleb cycle are insignificant. (A,B) Time-evolution of the cell centroid for three different values of the extracellular stress relaxation time (in seconds) for high density matrix (A) and low density matrix (B). A short FENE-P stress relaxation time corresponds with a stiffer extracellular fluid. (A, inset) Ellipses represent the polymeric stress tensor **σ**_**p**_ at a time-point during early bleb retraction. The major axis is aligned with the principal eigenvector of **σ**_**p**_, with length scaled on the associated eigenvalue. The minor axis is associated with the second eigenvector of **σ**_**p**_. The ellipses represent the directions and distension of the polymer field. Elastic stresses build up at the cell front and resist pressure-driven bleb expansion. Unless otherwise stated, polymeric model parameter values: 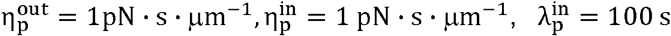. (C) Time-evolution of the cell centroid for three different values of the intracellular stress relaxation time (in seconds). Polymeric model parameter values: 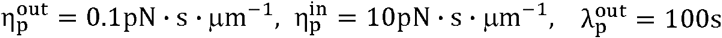. (D) Mean-square distention of the immersed polymer material tr(**σ**_**p**_) (blue-yellow scale bar) and local membrane elastic energy (gray-black scale bar) at a time-point during bleb retraction. Regions of high polymer stress and membrane elastic tension buildup at the cell front. (E–I) Time-evolution of the cell centroid for three different values of the number of myosin molecules inside the cell N_myo_ (E), for four different values of the cortical stiffness per unit of actin parameter κ_c_ (F), for different values of the fraction of the cell membrane perimeter (bleb size) ϕ_b_ that loses mechanical connection with the underlying cortex (G), for different values of the cortex-cytoplasm viscous drag coefficient γ_c_(pN· s · μm^−1^ (H), and for different values of the plasma membrane spring stiffness κ_m_(pN· μm^−1^ ) (I). High cortical contractility, soft cortex, intermediate bleb sizes, high cortex porosities and low plasma membrane stiffness maximize cell displacements during bleb expansion. In all panels, membrane-cortex linkers are broken by hand locally in a region set by ϕ_b_ at time t = 0. Cortex turnover parameter value: K_τ_ = 100.

**Figure S5.**
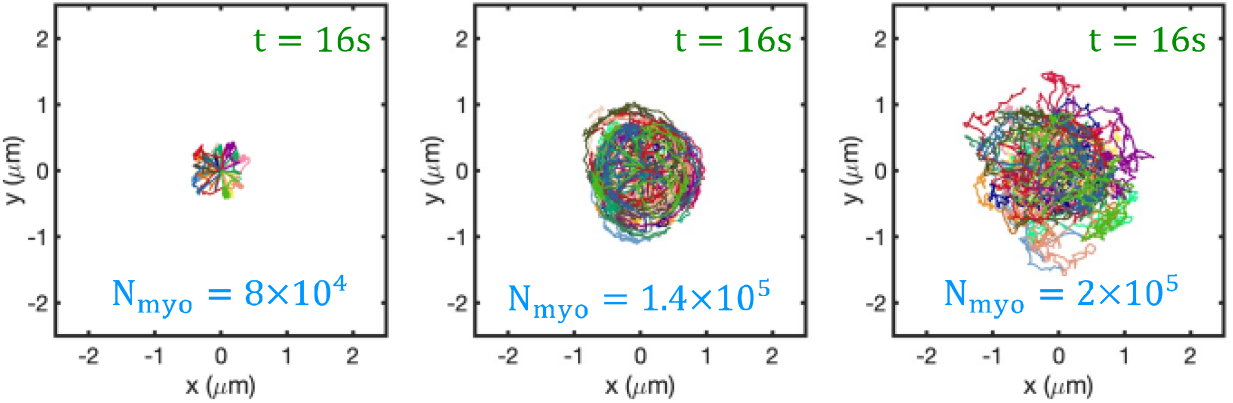
High contractile bleb-producing cells swim more efficiently. Wind rose plots depicting the migratory tracks of cells for three different myosin levels: low (left), intermediate (center), high (right). Each color represents a single-cell trajectory. Intermediate myosin levels (N_myo_ =1.4× 10^5^) build up a hydrostatic intracellular pressure of ∼50 pN/μm (see Fig. S1B), comparable to that of T cells under DMSO conditions ^21^.

**Figure S6.**
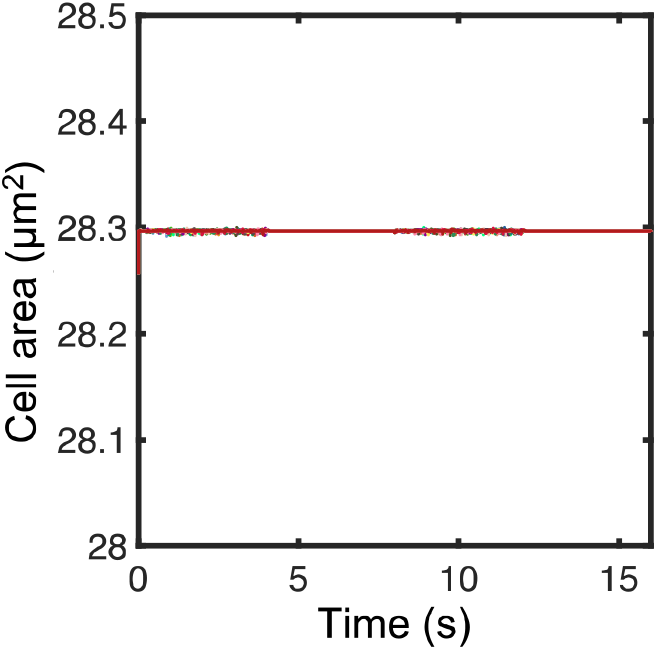
Time evolution of cell area for all the cells in Fig. S5 (center). Throughout the simulations, cell area remains nearly conserved, with only minor fluctuations arising from small water fluxes across the plasma membrane.

**Figure S7.**
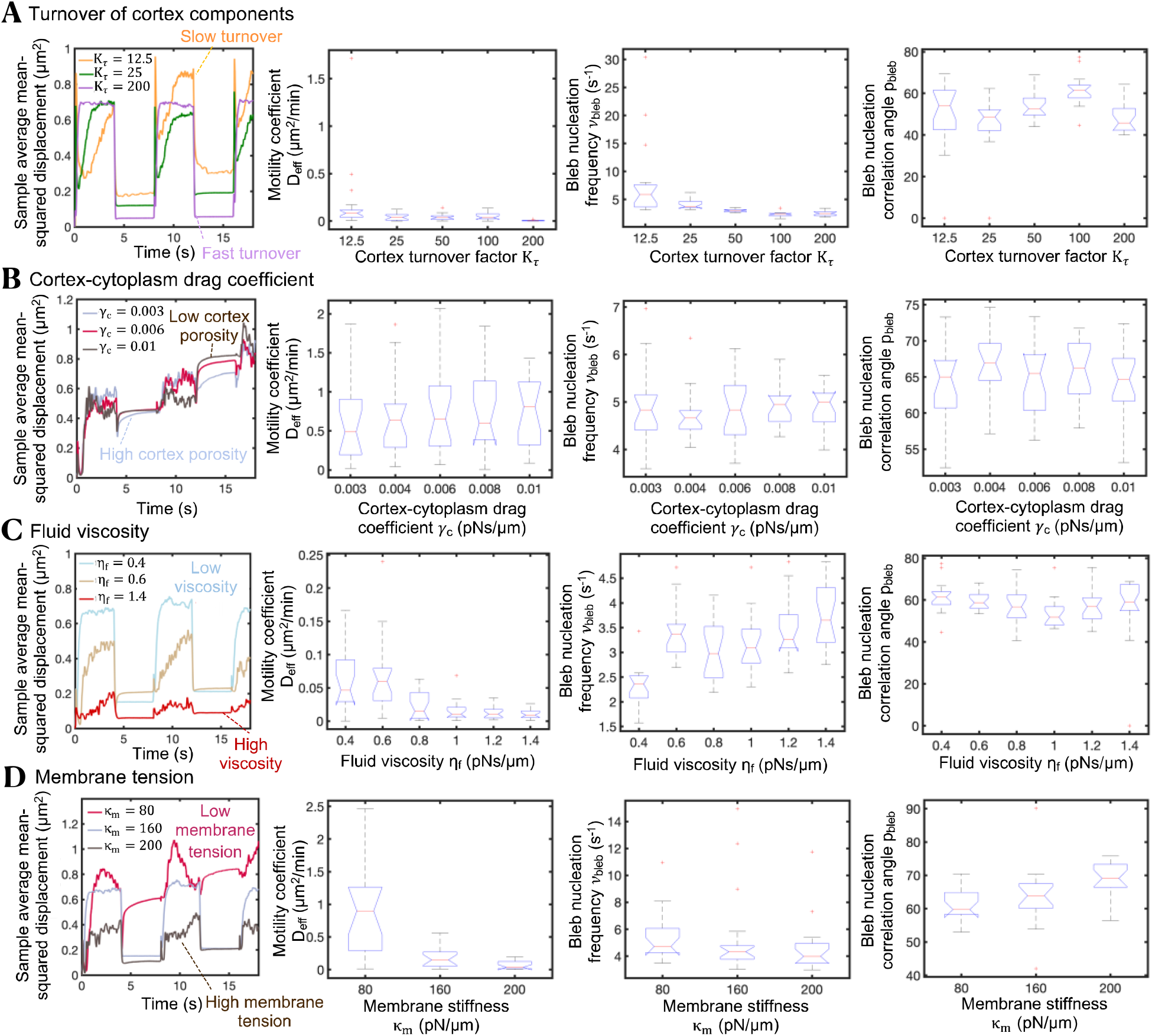
Cortex turnover, cortex porosity, fluid viscosity and membrane tension modulate bleb-based cell swimming capabilities. (A–D) Sample average mean-squared displacement, random motility coefficient D_eff_, bleb nucleation frequency *ν*_bleb_, and polar angle between consecutive bleb nucleation events p_bleb_ for different values of the (A) cortex turnover factor K_τ_ (B) cortex-cytoplasm drag coefficient γ (pN· s ·μm^−1^, (C) fluid viscosity η_f_ (pN· s · μm^−1^ ) and (D) membrane spring stiffness parameter κ_m_ (pN· μm^−1^ ).

**Figure S8.**
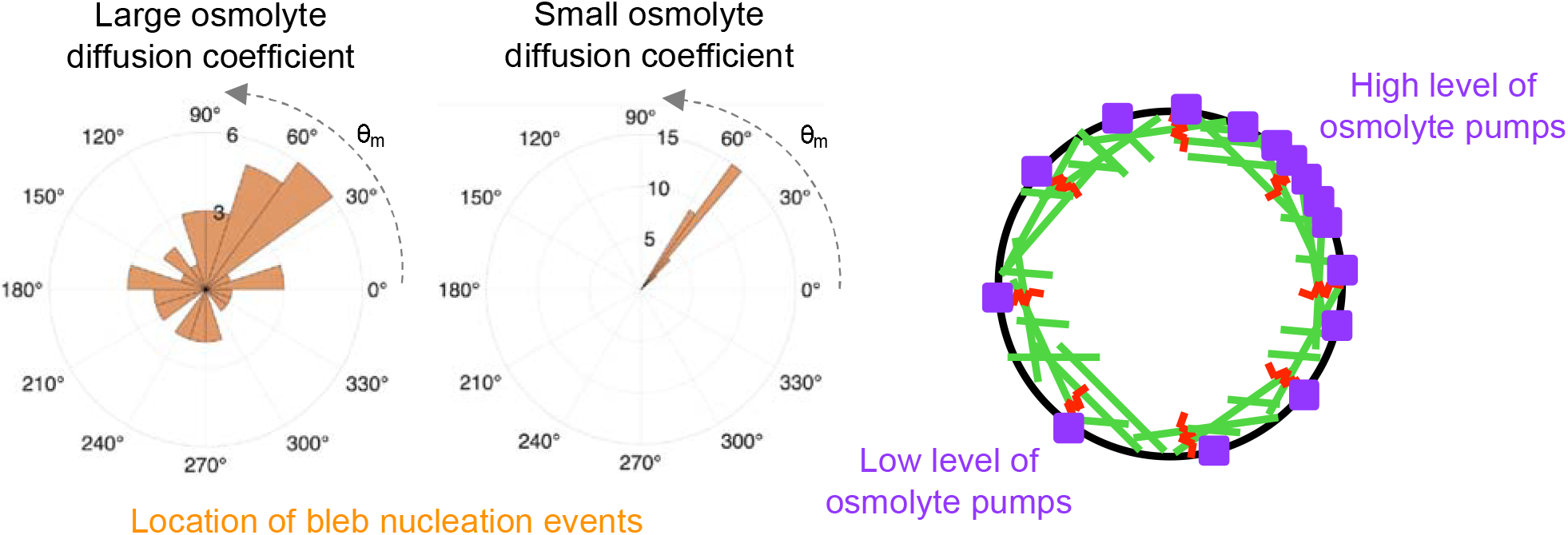
Blebs nucleate in high osmotic pressure regions. (A) Polar distribution of bleb nucleation events for fast osmolyte diffusion (D_in_ = 500µm^2^ /s) and slow intracellular osmolyte diffusion (D_in_ = 5µm^2^ /s), for an asymmetric osmolyte active pumping influx 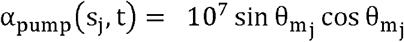, where 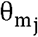 is the polar angle associated to the jth membrane node. Parameter values: D_out_ = 2D_in_ and α_passive_ = 10^−3^).

**Figure S9.**
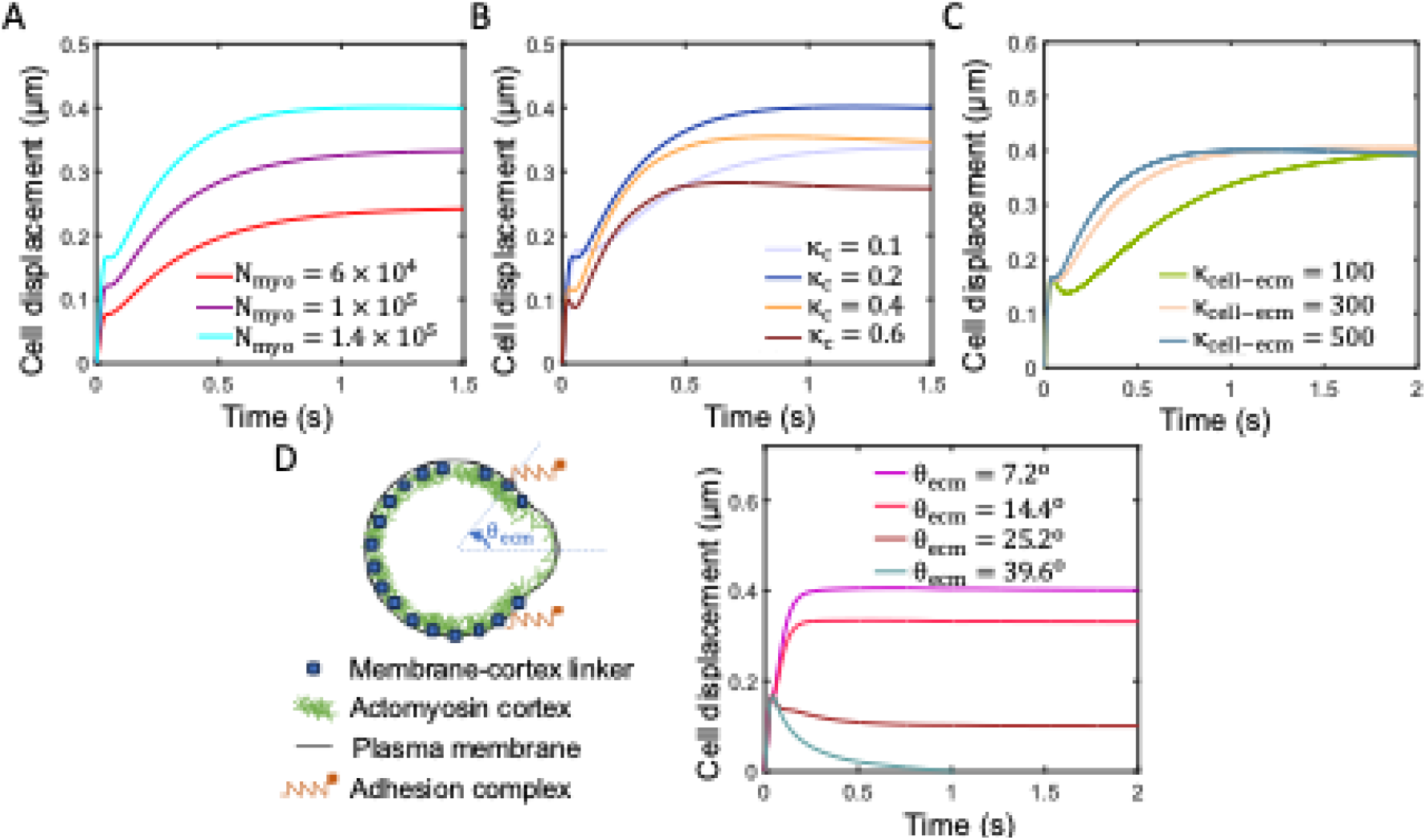
Substrate stiffness does not impact cell displacements in the hybrid adhesion-bleb migration mode. (A–C) Time-evolution of the cell centroid for three different values of the number of myosin molecules inside the cell N_myo_ (A), four different values of the cortical stiffness per unit of actin parameter κ_c_ (B), and for three different values of the effective stiffness of the cell-extracellular matrix tandem κ_cell–ecm_ (C). Larger cell displacements are achieved for elevated cortical tensions and intermediate cortical stiffnesses levels. (D) Time-evolution of the cell centroid for four different configurations of cell-matrix adhesion complex formation. Greater cell displacements are achieved when adhesions form at the cell’s front. Notice that the inclusion of two focal adhesions can mimic the squeezing of cells through narrow pores.

**Movie S1:** Fluid field (left), hydrostatic pressure (middle) and temporal evolution of cell displacements (right) during a single isolated bleb cycle in the absence of adhesion-based forces. Velocity field is expressed in μm/s and hydrostatic pressure in pN/μm.

**Movie S2:** Representative migration dynamics of a cell in the absence of cortical oscillations. Colors and vectors represent the magnitude and directionality of the fluid velocity (in μm/s .

**Movie S3:** Cortex and plasma membrane dynamics (left) and temporal evolution of cell displacements (right) during a single isolated bleb cycle in the presence of adhesion-based forces.

**Movie S4**: T cell migration in 3D LLS. Left panels: T cell migration in collagen I–conjugated LLS. Right panels: T cells from the same batch cultured in inert (unconjugated) LLS microgels. Two technical replicates (S1 and S2) are shown for each condition. Collagen-conjugated LLS microgels were labeled with Rhodamine B (red), and cells were labeled with CellTracker™ Green CMFDA.

**Movie S5**: A single T cell (green) migrating within the 3D interstitial space of collagen-conjugated LLS microgels (red).

**Movie S6**: A single T cell (green) migrating within the 3D interstitial space of inert (unconjugated) LLS microgels (red).

## Notes

### Competing Interest Statement

The authors have declared no competing interest.

### Summary of Updates

A few revisions have been made to ensure that conclusions remain consistent with computational and experimental results and existing literature

