## Supplemental text for "Biophysical modeling identifies an optimal hybrid amoeboid-mesenchymal mechanism for maximal T cell migration speeds"

**SUPPLEMENTAL INFORMATION**

**Model description**

We aim to investigate the cell migration capabilities of bleb-producing cells in an unbounded viscoelastic medium through the development of a two-dimensional biophysical mechanistic model. While the model would ideally be a three-dimensional model, we found that such a model is computationally too demanding for available supercomputing resources. Even so, the fundamental physics is captured, which has the potential to be recast into three dimensions in the future. The cell is composed of two distinct structures: a plasma membrane that defines the bounds of the cell and an actomyosin cortex that underlies the plasma membrane. Both are considered Lagrangian structures, and they are initially discretized as a linear chain of $N_{b}$ Lagrangian points. We follow their temporal trajectories through their vector positions $\mathbf{X}_{\mathbf{m}}\left( s_{j},t \right)$ and $\mathbf{X}_{\mathbf{c}}\left( s_{j},t \right)$, where the subscripts $m$ and $c$ denote membrane and cortex, respectively, and $s_{j}$ denotes the arc-length parameter associated to the j-th Lagrangian point. As the cell migrates through the environment, both membrane and cortex structures transmit forces on the surrounding medium, generating viscoelastic stresses. At the continuum level, conservation of mass and momentum on the viscoelastic fluid reads

$\nabla\cdot\mathbf{v}_{f}\left( \mathbf{x},t \right)=0 (S1)$

$\nabla\cdot\boldsymbol{\sigma}_{\mathrm{tot}}\left( \mathbf{x},t \right)+\mathcal{F}_{\mathrm{tot}}\left( \mathbf{x},t \right)=\mathbf{0} (S2)$

$\mathcal{F}_{\mathrm{tot}}\left( \mathbf{x},t \right)=\mathcal{F}_{\mathbf{m}}\left( \mathbf{x},t \right)+\mathcal{F}_{\mathbf{c, drag}}\left( \mathbf{x},t \right)+\mathcal{F}_{\mathbf{c,ecm}}\left( \mathbf{x},t \right) (S3)$

where $\mathbf{v}_{f}$ is the fluid velocity, $\boldsymbol{\sigma}_{\mathbf{tot}}$ is the total stress, $\mathcal{F}_{\mathbf{tot}}$ is the total force density (force per unit area) on the fluid, $\mathcal{F}_{\mathbf{m}}$ is the membrane force density, $\mathcal{F}_{\mathbf{c,ecm}}$ is the cell-matrix adhesion force density, and $\mathcal{F}_{\mathbf{c, drag}}$ is the cortical force density generated in the intracellular medium. Notice that the cell-matrix adhesion force has only been included in the model to generate hybrid bleb-based adhesion-based cell migration results shown in Fig. 6. The whole domain $\Omega$ is discretized with a spatially uniform rectangular grid with $N$ nodes in both $x$ and $y$ directions. The variables $\mathbf{v}_{f}$, $\boldsymbol{\sigma}_{\mathrm{tot}}$ and $\mathcal{F}_{\mathrm{tot}}$ are defined in this Eulerian grid over the whole domain. Notice that inertial effects for the immersed elastic structures and viscoelastic fluid are negligible. The lack of inertia together with Newton’s laws of motion implies that the sum of all the forces that the cell exerts on the surrounding medium always vanish: $\int_{\Omega} \mathcal{F}_{\mathrm{tot}}\left( \mathbf{x},t \right)d\mathbf{x}=\mathbf{0}$, consistent with Eq. (S2). By construction, this condition is always satisfied, ensuring that the model obeys conservation of momentum at all times. The total stress $\boldsymbol{\sigma}_{\mathbf{tot}}\boldsymbol{=}\boldsymbol{\sigma}_{\mathbf{f}}\boldsymbol{+}\boldsymbol{\sigma}_{\mathbf{p}}$ is the sum of two contributions: a purely viscous stress $\boldsymbol{\sigma}_{\mathbf{f}}$ and an extra polymeric viscoelastic stress $\boldsymbol{\sigma}_{\mathbf{p}}$:

$$\boldsymbol{\sigma}_{\mathbf{f}}=-\boldsymbol{\nabla}p+\eta_{f}\left( \boldsymbol{\nabla v}_{\mathbf{f}}+{\boldsymbol{\nabla v}_{\mathbf{f}}}^{\mathbf{T}} \right) (S4)$$

$$\boldsymbol{\sigma}_{\mathbf{p}}=\frac{\eta_{p}(\mathbf{x},t)}{\lambda_{p}(\mathbf{x},t)}\left[ \left( \frac{\boldsymbol{\kappa}_{\mathbf{p}}}{1-{\mathrm{tr}\left( \boldsymbol{\kappa}_{\mathbf{p}} \right)}/{L_{p}^{2}}} \right)-\mathbf{I} \right] (S5)$$

where we have chosen the FENE-P constitutive relation for the stress to model the viscoelastic nature of the intracellular and extracellular spaces^1^. The original FENE-P model was developed by Bird and coworkers^2^, and a few variations of their model have been used since then^3^. In our current study, we have used the FENE-P constitutive equation of Housiadas & Beris^1^ to model the cellular cytoplasm and cell surroundings. In Eqs. (S4) and (S5), $p$ is the hydrostatic pressure, $\eta_{f}$ and $\eta_{p}$ are, respectively, the fluid viscosity and polymer viscosity, $\lambda_{p}$ is the polymer stress relaxation time, $\boldsymbol{\kappa}_{\mathbf{p}}$ is the conformation stress tensor and $L_{p}$ is the polymer extensibility parameter, a measure of the maximum polymeric deformation. The intracellular and extracellular spaces are considered viscoelastic fluids with different rheological properties. We denote the intracellular and extracellular polymer viscosities as $\eta_{p}^{\mathrm{in}}$ and $\eta_{p}^{\mathrm{out}}$, and the intracellular and extracellular stress relaxation times as $\lambda_{p}^{\mathrm{in}}$ and $\lambda_{p}^{\mathrm{out}}$, respectively. As a guidance, the FENE-P viscoelastic model approximates the cellular environment as a suspension of entropic dumbbells with density $n_{\mathrm{dum}}$. Each dumbbell can be thought of as two spheres linked by an elastic spring with stiffness $\kappa_{\mathrm{dum}}$. As the spheres that comprise each dumbbell move through the medium, they are subjected to drag forces, with drag coefficient $\mu_{\mathrm{dum}}$. The FENE-P macroscopic properties polymer viscosity $\eta_{p}$ and polymer stress relaxation time $\lambda_{p}$ can then be related to the dumbbell microscopic properties and scale as $\eta_{p}\propto n_{\mathrm{dum}}\mu_{\mathrm{dum}}$ and $\lambda_{p}\propto\mu_{\mathrm{dum}}/\kappa_{\mathrm{dum}}$, respectively. In order to solve the problem numerically, we define a continuous polymer viscosity and stress relaxation time in the whole domain following a similar procedure to that used in a prior numerical study^4^:

$$\eta_{p}\left( \mathbf{x},t \right)=\eta_{p}^{\mathrm{in}}+\left( \eta_{p}^{\mathrm{out}}-\eta_{p}^{\mathrm{in}} \right)H\left( d\left( \mathbf{x} \right) \right), \lambda_{p}\left( \mathbf{x},t \right)=\lambda_{p}^{\mathrm{in}}+\left( \lambda_{p}^{\mathrm{out}}-\lambda_{p}^{\mathrm{in}} \right)H\left( d\left( \mathbf{x} \right) \right) (S6-S7)$$

where $H\left( d \right)$ is the discrete Heaviside function:

$$H\left( d \right)=\left\{ \begin{aligned} 0 d<-2h \\ \frac{1}{2}\left( 1+\frac{d}{2h}+\frac{1}{\pi}\sin\left( \frac{\pi d}{2h} \right) \right) -2h\leq d\leq2h \\ 1 d<2h \end{aligned} (S8) \right.$$

d$\left( \mathbf{x} \right)$ is the shortest distance to the cell membrane from the point $\mathbf{x}$, and $h$ is the grid size of the Eulerian grid. The time evolution of the conformation stress tensor $\boldsymbol{\kappa}_{\mathbf{p}}$ reads

$$\frac{\partial\boldsymbol{\kappa}_{\mathbf{p}}}{\partial t}=\frac{1}{\lambda_{p}}\left( \mathbf{I}-\frac{\boldsymbol{\kappa}_{\mathbf{p}}}{1-{\mathrm{tr}\left( \boldsymbol{\kappa}_{\mathbf{p}} \right)}/{L_{p}^{2}}} \right)-\mathbf{v}_{\mathbf{f}}\boldsymbol{\cdot}\boldsymbol{\nabla}\boldsymbol{\kappa}_{\mathbf{p}}\boldsymbol{+}{\boldsymbol{\nabla v}_{\mathbf{f}}}^{\mathbf{T}}\boldsymbol{\cdot}\boldsymbol{\kappa}_{\mathbf{p}}\boldsymbol{+}\boldsymbol{\kappa}_{\mathbf{p}}\boldsymbol{\cdot}\boldsymbol{\nabla v}_{\mathbf{f}} (S9)$$

The first term on the right-hand side (RHS) of Eq. (S9) captures polymer stress relaxation kinetics, the second term captures polymer advection, and the last two terms capture rotation and deformation of the polymeric material.

The cell membrane is an elastic structure subjected to tension, bending, membrane-cortex adhesion and short-range repulsive forces. The total force per unit length on the cell membrane $\mathbf{F}_{\mathbf{m}}$ is thus

$$\mathbf{F}_{\mathbf{m}}\left( s,t \right)=\mathbf{F}_{\mathbf{m,tens}}\left( s,t \right)+\mathbf{F}_{\mathbf{m,bend}}\left( s,t \right)+\mathbf{F}_{\mathbf{m,adh}}\left( s,t \right)+\mathbf{F}_{\mathbf{m,rep}}\left( s,t \right) (S10)$$

We assume a linear stress-strain relation for membrane tension^5^; accordingly the membrane tension force per unit length on the membrane element $j$ reads

$$\mathbf{F}_{\mathbf{m,tens}}\left( s_{j}\mathbf{,t} \right)=-F_{m_{0}}^{\mathrm{tens}}{\mathbf{n}_{\mathbf{m}}}_{\mathbf{j}}+\kappa_{m}\left[ \left( {d_{m}}_{j}-{\mathcal{l}_{m}}_{0} \right){\boldsymbol{\tau}_{\mathbf{m}}}_{\mathbf{j}}-\left( {d_{m}}_{j-1}-{\mathcal{l}_{m}}_{0} \right){\boldsymbol{\tau}_{\mathbf{m}}}_{\mathbf{j-1}} \right] (S11)$$

where $F_{m_{0}}^{\mathrm{tens}}$ is the resting membrane tension force, assumed to be spatially uniform, ${\mathbf{n}_{\mathbf{m}}}_{\mathbf{j}}$ is the membrane outwards unit normal, $\kappa_{m}$ is the plasma membrane spring stiffness parameter, ${d_{m}}_{j}=\left| {\mathbf{X}_{\mathbf{m}}}_{\mathbf{j+1}}\mathbf{-}{\mathbf{X}_{\mathbf{m}}}_{\mathbf{j}} \right|$ is the distance between membrane neighbor elements, ${\boldsymbol{\tau}_{\mathbf{m}}}_{\mathbf{j}}\boldsymbol{=}\left( {\mathbf{X}_{\mathbf{m}}}_{\mathbf{j+1}}\mathbf{-}{\mathbf{X}_{\mathbf{m}}}_{\mathbf{j}} \right)\boldsymbol{/}\left| {\mathbf{X}_{\mathbf{m}}}_{\mathbf{j+1}}\mathbf{-}{\mathbf{X}_{\mathbf{m}}}_{\mathbf{j}} \right|$ is the membrane tangent unit vector, and ${\mathcal{l}_{m}}_{0}$ is the membrane spring resting length, which corresponds with the initial separation distance between neighbor membrane element points. We consider a simple quadratic dependence of the elastic bending energy on membrane curvature, thus the membrane bending force per unit length reads^6^

$$\mathbf{F}_{\mathbf{m,bend}}\left( s,t \right)=\beta_{m}\frac{\partial^{4}\mathbf{X}_{\mathbf{m}}}{\partial s^{4}}, (S12)$$

where $\beta_{m}$ is the membrane bending stiffness. The membrane-cortex adhesion energy is assumed to be quadratic with respect to the membrane-cortex linker deformation; the membrane-cortex adhesion force per unit length on the membrane is thus given by

$$\mathbf{F}_{\mathbf{m,adh}}\left( s_{j},t \right)=-\rho_{\mathrm{adh}}\left( s_{j},t \right)\kappa_{\mathrm{adh}}\left( {d_{\mathrm{adh}}}_{j}-{\mathcal{l}_{\mathrm{adh}}}_{0} \right){\boldsymbol{\tau}_{\mathbf{adh}}}_{\mathbf{j}} (S13)$$

where we have modeled membrane-cortex elastic linkers as linear elastic springs. Here, $\rho_{\mathrm{adh}}$ is the local density of membrane-cortex linkers engaged on each membrane-cortex connection, $\kappa_{\mathrm{adh}}$ is the elastic stiffness of each membrane-cortex linker, $d_{\mathrm{adh}}=\left| {\mathbf{X}_{\mathbf{m}}}_{\mathbf{j}}\mathbf{-}{\mathbf{X}_{\mathbf{c}}}_{\mathbf{j}} \right|$ and ${\mathcal{l}_{\mathrm{adh}}}_{0}$ are the stretching and resting length of linkers, respectively, and ${\boldsymbol{\tau}_{\mathbf{adh}}}_{\mathbf{j}}\boldsymbol{=}\left( {\mathbf{X}_{\mathbf{m}}}_{\mathbf{j}}\mathbf{-}{\mathbf{X}_{\mathbf{c}}}_{\mathbf{j}} \right)\boldsymbol{/}\left| {\mathbf{X}_{\mathbf{m}}}_{\mathbf{j}}\mathbf{-}{\mathbf{X}_{\mathbf{c}}}_{\mathbf{j}} \right|$ is a unit vector whose direction is set by the relative position of mechanically linked membrane-cortex elements. The stability of membrane-cortex linkers strongly depends on active forces and membrane-cortex adhesion properties. We follow two distinct approaches to capture the kinetics of membrane-cortex linkers: a deterministic approach and a stochastic approach.

Deterministic model: To elucidate cell migration dynamics during a single bleb migration cycle we initially break membrane-cortex adhesion linkers by hand on a local cellular region, whose size is given by $\phi_{b}$, the fraction of the plasma membrane perimeter (neck bleb size) that loses mechanical connection with the underlying cortex. Membrane-cortex adhesion loss is followed by recruitment of new cortex. Mass conservation of membrane-cortex linkers follows simple association and dissociation kinetics:

$$\frac{\partial n_{\mathrm{adh}}\left( s_{j},t \right)}{\partial t}=k_{\mathrm{adh}}^{\mathrm{on}}\rho_{\mathrm{adh}}^{\mathrm{free}}-k_{\mathrm{adh}}^{\mathrm{off}} n_{\mathrm{adh}}\left( s_{j},t \right)\boldsymbol{,}(S14)$$

where $n_{\mathrm{adh}}=\int_{s_{j-1/2}}^{s_{j+1/2}} \rho_{\mathrm{adh}}\mathrm{ds}$ is the discretized effective number of membrane-cortex linkers at position $s_{j}$ at time $t$, $k_{\mathrm{adh}}^{\mathrm{on}}$ is the force-independent linker association rate constant and $k_{\mathrm{adh}}^{\mathrm{off}}$ is the force-dependent linker dissociation rate constant. The concentration of free linkers in the cytoplasm has been denoted by $\rho_{\mathrm{adh}}^{\mathrm{free}}=\left( N_{\mathrm{adh}}^{\mathrm{tot}}-\sum_{k=1}^{N_{b}} n_{\mathrm{adh}}\left( s_{k},t \right) \right)/{A_{\mathrm{cell}}h_{\mathrm{cell}}}$, where $N_{\mathrm{adh}}^{\mathrm{tot}}$ is the total number of linkers available in the cell and $h_{\mathrm{cell}}$ is the cell thickness in the z-direction.

Stochastic model: To account for changes in the dynamics of cortex component amounts we introduce the stochastic cell migration model counterpart. In this stochastic version, we model membrane-cortex linker kinetics as jump processes of unit size that follow Poisson statistics. Membrane-cortex linkers are not broken by hand. Instead, the stability of membrane-cortex linkers strongly depends on active forces and membrane-cortex adhesion properties. Membrane-cortex linkers stochastically associate at a force-independent rate $k_{\mathrm{adh}}^{\mathrm{on}}\rho_{\mathrm{adh}}^{\mathrm{free}}$ and unbind by force with an effective dissociation rate that increases exponentially with force according to Bell’s law^7^: $k_{\mathrm{adh}}^{\mathrm{off}}e^{k_{\mathrm{adh}}\left( \left| {d_{\mathrm{adh}}}_{j}-{\mathcal{l}_{\mathrm{adh}}}_{0} \right|/{F_{\mathrm{adh}}^{\mathrm{rupt}}} \right)}$, where the linker unloaded dissociation rate and the characteristic linker rupture force have been denoted as $k_{\mathrm{adh}}^{\mathrm{off}}$ and $F_{\mathrm{adh}}^{\mathrm{rupt}}$, respectively. The probability that a membrane-cortex linker associates on a given membrane/cortical element in an interval of time $\Delta t$ is $p_{\mathrm{adh}}^{\mathrm{on}}=1-e^{-k_{\mathrm{adh}}^{\mathrm{on}}\rho_{\mathrm{adh}}^{\mathrm{free}}\Delta t}$. For a small enough timestep $(p_{\mathrm{adh}}^{\mathrm{on}}\ll1)$, $p_{\mathrm{adh}}^{\mathrm{on}}\approx k_{\mathrm{adh}}^{\mathrm{on}}\rho_{\mathrm{adh}}^{\mathrm{free}}\Delta t$. Similarly, the probability that a membrane-cortex linker dissociates from a given membrane/cortical element in an interval of time $\Delta t$ is $p_{\mathrm{adh}}^{\mathrm{off}}\approx k_{\mathrm{adh}}^{\mathrm{off}}e^{k_{\mathrm{adh}}\left( \left| {d_{\mathrm{adh}}}_{j}-{\mathcal{l}_{\mathrm{adh}}}_{0} \right|/{F_{\mathrm{adh}}^{\mathrm{rupt}}} \right)}\Delta t$. At each time step, number of linkers associated to each cortical element is updated by generating a uniformly-distributed random number between 0 and 1, and comparing the generated random number to the probability of the process under consideration. If the random number is less than the probability, then it is assumed that the event occurred, and the system variables are updated accordingly.

To prevent unphysical membrane-cortex crossings, we introduce a short-range membrane-cortex repulsive force. The repulsive force on the j-th membrane node is given by:

$$\mathbf{F}_{\mathbf{m,rep}}\left( s_{j},t \right)=\left\{ \begin{aligned} \kappa_{\mathrm{rep}}\sum_{i} \left| {d_{\mathrm{adh}}}_{\mathrm{ji}}-\mathcal{l}_{\mathrm{rep}} \right|{\boldsymbol{\tau}_{\mathbf{adh}}}_{\mathbf{ji}} \\ 0 \end{aligned} \right.\begin{matrix} \mathrm{if}{d_{\mathrm{adh}}}_{\mathrm{ji}}<\mathcal{l}_{\mathrm{rep}} \\ \mathrm{if} {d_{\mathrm{adh}}}_{\mathrm{ji}}>\mathcal{l}_{\mathrm{rep}} \end{matrix} (S15)$$

where ${d_{\mathrm{adh}}}_{\mathrm{ji}}$ is the distance between the j-th membrane node and the i-th cortical node, ${\boldsymbol{\tau}_{\mathbf{adh}}}_{\mathbf{ji}}\boldsymbol{=}\left( {\mathbf{X}_{\mathbf{m}}}_{\mathbf{j}}\mathbf{-}{\mathbf{X}_{\mathbf{c}}}_{\mathbf{i}} \right)\boldsymbol{/}\left| {\mathbf{X}_{\mathbf{m}}}_{\mathbf{j}}\mathbf{-}{\mathbf{X}_{\mathbf{c}}}_{\mathbf{i}} \right|$, and $\mathcal{l}_{\mathrm{rep}}$ is the short-range repulsive cutoff distance.

The cortex is considered a cross-linked actomyosin poroelastic structure that generates contractile forces creating tension in the cortical network. It is initially modeled as a linear chain of beads jointed by elastic Hookean springs, that are additionally adhered to the plasma membrane by the linear elastic linkers mentioned above. Notice that upon complete membrane-cortex adhesion loss ($\rho_{\mathrm{adh}}\left( s_{j},t \right)=0$), after a refractory time $t_{\mathrm{refr}}$, a new cortical element appears underneath the cell membrane under stress-free conditions to initiate new cortex formation, and the old cortical element components are transferred into the cytoplasm. Force balance on each cortical element reads

$$\mathbf{f}_{\mathbf{c, elast}}\left( s,t \right)+\mathbf{f}_{\mathbf{c, myo}}\left( s,t \right)+\mathbf{f}_{\mathbf{c, adh}}\left( s,t \right)+\mathbf{f}_{\mathbf{c, rep}}\left( s,t \right)+\mathbf{f}_{\mathbf{c, ecm}}\left( s,t \right)+\mathbf{f}_{\mathbf{c, drag}}\left( s,t \right)=\boldsymbol{0} (S16)$$

The elastic cortical force $\mathbf{f}_{\mathbf{c, elast}}$ is associated with the tensional state of the cortex and it is given by

$$\mathbf{f}_{\mathbf{c, elast}}\left( s_{j},t \right)=\frac{1}{2}\left( \kappa_{c_{j+1}}^{\mathrm{eff}}+\kappa_{c_{j}}^{\mathrm{eff}} \right)\left[ \left( {d_{c}}_{j}-{\mathcal{l}_{c}}_{0} \right){\boldsymbol{\tau}_{\mathbf{c}}}_{\mathbf{j}} \right]-\frac{1}{2}\left( \kappa_{c_{j}}^{\mathrm{eff}}+\kappa_{c_{j-1}}^{\mathrm{eff}} \right)\left[ \left( {d_{c}}_{j-1}-{\mathcal{l}_{c}}_{0} \right){\boldsymbol{\tau}_{\mathbf{c}}}_{\mathbf{j-1}} \right] (S17)$$

where $\kappa_{c_{j}}^{\mathrm{eff}}$ is the effective cortex spring stiffness at the cortical node $j$, ${d_{c}}_{j}=\left| {\mathbf{X}_{\mathbf{c}}}_{\mathbf{j+1}}\mathbf{-}{\mathbf{X}_{\mathbf{c}}}_{\mathbf{j}} \right|$ is the distance between cortical neighbor elements, ${\boldsymbol{\tau}_{\mathbf{c}}}_{\mathbf{j}}\boldsymbol{=}\left( {\mathbf{X}_{\mathbf{c}}}_{\mathbf{j+1}}\mathbf{-}{\mathbf{X}_{\mathbf{c}}}_{\mathbf{j}} \right)\boldsymbol{/}\left| {\mathbf{X}_{\mathbf{c}}}_{\mathbf{j+1}}\mathbf{-}{\mathbf{X}_{\mathbf{c}}}_{\mathbf{j}} \right|$ is the cortical tangent unit vector, and ${\mathcal{l}_{c}}_{0}$ is the cortex spring resting length, which corresponds with the initial separation distance between neighbor cortical element points. The effective cortex spring stiffness is assumed to depend proportionally to the amount of local actin density as $\kappa_{c}^{\mathrm{eff}}\left( s,t \right)=\kappa_{c}n_{\mathrm{act}}\left( s,t \right)$, where $\kappa_{c}$ is the cortex stiffness per unit of actin. The myosin-mediated cortical tension forces $\mathbf{f}_{\mathbf{c, myo}}$ obey a linear force-velocity relationship, which captures how the speed of actin filaments depends on the applied force. At low opposing force, myosin motors cycle rapidly, leading to high contraction velocity. As the opposing force increases, myosin heads spend more time in force-generating states, and contraction velocity slows down eventually to stall. Mathematically, this relationship is expressed as^8,9^

$$\mathbf{f}_{\mathbf{c, myo}}\left( s_{j},t \right)\mathbf{=}\frac{1}{2}\left( {F_{st,myo}^{\mathrm{eff}}}_{j+1}+{F_{st,myo}^{\mathrm{eff}}}_{j} \right)\left[ {\boldsymbol{\tau}_{\mathbf{c}}}_{\mathbf{j}}-\frac{\left( {\mathbf{V}_{\mathbf{c}}}_{\mathbf{j}}-{\mathbf{V}_{\mathbf{c}}}_{\mathbf{j+1}} \right)}{v_{0}^{\mathrm{myo}}}\cdot{\boldsymbol{\tau}_{\mathbf{c}}}_{\mathbf{j}}{\boldsymbol{\tau}_{\mathbf{c}}}_{\mathbf{j}} \right]+\frac{1}{2}\left( {F_{st,myo}^{\mathrm{eff}}}_{j}+{F_{st,myo}^{\mathrm{eff}}}_{j-1} \right)\left[ -{\boldsymbol{\tau}_{\mathbf{c}}}_{\mathbf{j-1}}-\frac{\left( {\mathbf{V}_{\mathbf{c}}}_{\mathbf{j}}-{\mathbf{V}_{\mathbf{c}}}_{\mathbf{j-1}} \right)}{v_{0}^{\mathrm{myo}}}\cdot{\boldsymbol{\tau}_{\mathbf{c}}}_{\mathbf{j-1}}{\boldsymbol{\tau}_{\mathbf{c}}}_{\mathbf{j-1}} \right] (S18)$$

where ${F_{st,myo}^{\mathrm{eff}}}_{j}$ is the effective myosin stall force at cortical node $j$, ${\mathbf{V}_{\mathbf{c}}}_{\mathbf{j}}$ is the cortex velocity vector of the j-th node, and $v_{0}^{\mathrm{myo}}$ is the load-free myosin velocity. Notice that the actomyosin force has been projected along the line joining the linked cortical nodes. The effective myosin stall force is assumed to depend proportionally to local myosin and actin amounts as $F_{st,myo}^{\mathrm{eff}}\left( s,t \right)=F_{\mathrm{st}}S_{\mathrm{osc}}n_{\mathrm{myo}}\left( s,t \right)n_{\mathrm{act}}\left( s,t \right)$, where $F_{\mathrm{st}}$ is the myosin stall force per unit of myosin and actin, and $S_{\mathrm{osc}}$ is a temporal oscillatory signal that mimics periods of high and low cortical tensions. Notice that F_st is the critical model parameter responsible for the generation of cortical tension in the model.

Since cortical forces get transmitted to the plasma membrane through membrane-cortex linkers, they generate a buildup of intracellular hydrostatic pressure. Once the transmission of this squeezing actomyosin force to the plasma membrane is interrupted (due to mechanical disengagement between the membrane and cortex), the local intracellular hydrostatic pressure decreases. Essentially, the jump in hydrostatic pressure between intracellular and extracellular spaces results from in-plane forces on the plasma membrane. We assume that $S_{\mathrm{osc}}$ is a two-level periodic square wave of period $T_{\mathrm{myo}}$, where $S_{\mathrm{osc}}=1$ during high cortical tension and $S_{\mathrm{osc}}=0.1$ during low cortical tension. In Eqs. (S17) and (S18), the effective cortex spring stiffness and myosin stall force for each two-node cortical connection has been taken to be the average of the spring stiffness and stall force of the two nodes, respectively. This practice ensures that linear momentum is conserved in our model. The membrane-cortex adhesion force acting on the cortex is equal and opposite to the membrane-cortex adhesion force acting on the membrane. Considering that membrane forces are defined per unit length in Eq. (S10), and that cortical forces are net forces in Eq. (S13), membrane-cortex adhesion forces acting on the cortex read $\mathbf{f}_{\mathbf{c}}^{\mathbf{adh}}\left( s_{j},t \right)=-\mathbf{F}_{\mathbf{m}}^{\mathbf{adh}}\left( s_{j},t \right)\left( s_{j+1}-s_{j-1} \right)/2$.

Adherent blebby cells can use a hybrid adhesion-based bleb-based mode of migration to move through tissues. To study migration capabilities of adherent blebby cells, we use the deterministic model to investigate the effect of focal adhesion formation on cell displacements during an isolated bleb cycle (Fig. 6). We assume that, following bleb expansion, the cell forms a focal adhesion at the cell front with the extracellular matrix, a highly crosslinked fibrous network embedded in a fluid. Localized cellular adhesion forces on the matrix will get transmitted to the entire fibrous network. However, for simplicity, we assume that the effective force on the matrix is represented as a point force, whose point of application is denoted by the matrix node vector position $\mathbf{X}_{\mathbf{ecm}}$. The cortex therefore is subject to an adhesion force during bleb retraction given by $\mathbf{f}_{\mathbf{c, ecm}}\left( s_{m},t \right)=-\kappa_{cell-ecm}\left[ \left( d_{c,ecm}-{\mathcal{l}_{c,ecm}}_{0} \right)\boldsymbol{\tau}_{\mathbf{c,ecm}} \right]$, where $\kappa_{cell-ecm}$ is the effective stiffness of the cell adhesion protein complex-extracellular matrix tandem, $d_{c,ecm}=\left| {\mathbf{X}_{\mathbf{c}}}_{\mathbf{ecm}}\mathbf{-}\mathbf{X}_{\mathbf{ecm}} \right|$ is the distance between the m-th cortical node, involved in the mechanical interaction with the matrix and represented by the vector position ${\mathbf{X}_{\mathbf{c}}}_{\mathbf{ecm}}\boldsymbol{,}$and the position of the matrix node, ${\mathcal{l}_{c,ecm}}_{0}$ is the spring resting length, and $\boldsymbol{\tau}_{\mathbf{c,ecm}}$ is the unit vector joining the cortical node and matrix node. We assume that the matrix is not compliant, i.e., the matrix node is stationary. Notice that we have only explored the hybrid bleb-adhesion-based cell migration mode during a single bleb cycle. The mechanism by which cells coordinate the spatial and temporal assembly and disassembly of focal adhesions, actin polymerization and traction forces in conjunction with blebbing are still to be elucidated.

The actin cortex also experiences drag forces as cytoplasmic material flows through the porous actomyosin network when membrane-cortex adhesion is lost; they are given by

$$\mathbf{f}_{\mathbf{c, drag}}\left( s_{j},t \right)=-\gamma_{c}\left( {\mathbf{V}_{\mathbf{c}}}_{\mathbf{j}}-\mathbf{v}_{f}\left( {\mathbf{X}_{\mathbf{c}}}_{\mathbf{j}},t \right) \right) (S19)$$

where $\gamma_{c}$ is the cortex-cytoplasm drag coefficient. The short-range membrane-cortex repulsive force on the cortex $\mathbf{f}_{\mathbf{c, rep}}\left( s,t \right)$ is equal and opposite to the repulsive force on the membrane (see Eq. (S15)).

Following a similar treatment to that of membrane-cortex linkers, we follow two distinct approaches to enforce mass conservation of actin and myosin: a deterministic approach and a stochastic approach.

Deterministic model: Conservation of cortical actin and myosin reads

$$\frac{\partial n_{\mathrm{act}}\left( s_{j},t \right)}{\partial t}=k_{\mathrm{act}}^{\mathrm{on}}\rho_{\mathrm{act}}^{\mathrm{free}}n_{\mathrm{adh}}\left( s_{j},t \right)-k_{\mathrm{act}}^{\mathrm{off}} n_{\mathrm{act}}\left( s_{j},t \right) (S20)$$

$$\frac{\partial n_{\mathrm{myo}}\left( s_{j},t \right)}{\partial t}=k_{\mathrm{myo}}^{\mathrm{on}}\rho_{\mathrm{myo}}^{\mathrm{free}}n_{\mathrm{act}}\left( s_{j},t \right)-k_{\mathrm{myo}}^{\mathrm{off}} n_{\mathrm{myo}}\left( s_{j},t \right) (S21)$$

where $k_{\mathrm{act}}^{\mathrm{on}}$ and $k_{\mathrm{act}}^{\mathrm{off}}$ are the actin association and dissociation constants and $k_{\mathrm{myo}}^{\mathrm{on}}$ and $k_{\mathrm{myo}}^{\mathrm{off}}$ are the myosin association and dissociation constants. The concentration of G-actin units in the cytoplasm has been denoted by $\rho_{\mathrm{act}}^{\mathrm{free}}=\left( N_{\mathrm{act}}^{\mathrm{tot}}-\sum_{k=1}^{N_{b}} n_{\mathrm{act}}\left( s_{k},t \right) \right)/{A_{\mathrm{cell}}h_{\mathrm{cell}}}$, where $N_{\mathrm{act}}^{\mathrm{tot}}$ is the total number of actin units available in the cell. Similarly, the concentration of myosin units in the cytoplasm has been denoted by $\rho_{\mathrm{myo}}^{\mathrm{free}}=\left( N_{\mathrm{myo}}^{\mathrm{tot}}-\sum_{k=1}^{N_{b}} n_{\mathrm{myo}}\left( s_{k},t \right) \right)/{A_{\mathrm{cell}}h_{\mathrm{cell}}}$, where $N_{\mathrm{myo}}^{\mathrm{tot}}$ is the total number of myosin units available in the cell. In Eqs. (S20) and (S21) we have assumed that recruitment of actin and myosin obeys first-order binding kinetics to linkers and actin, respectively. Upon a local membrane-cortex adhesion loss, the old cortical nodes are retained, we assume that actin and myosin do not associate to the local detached cortex; consequently, we set the association rate constants $k_{\mathrm{act}}^{\mathrm{on}}$ and $k_{\mathrm{myo}}^{\mathrm{on}}$ associated to the detached cortical node to 0 and actin and myosin eventually disappear at the old cortex element with rate constants $k_{\mathrm{act}}^{\mathrm{off}}$ and $k_{\mathrm{myo}}^{\mathrm{off}}$, respectively. When a new cortical node is added underneath of the cell membrane, actin and myosin are recruited at the node, allowing again the transmission of cortical forces to the cell membrane. We have assumed that the kinetics of cortex component amounts are governed by rapid association and dissociation kinetics, and transport of cortical components driven by cortical flows have been neglected.

Stochastic model: we model cortical kinetics as jump processes of unit size that follow Poisson statistics. The probabilities of an actin unit and myosin unit associating on a membrane/cortical element in an interval of time $\Delta t$ are $p_{\mathrm{act}}^{\mathrm{on}}\approx k_{\mathrm{act}}^{\mathrm{on}}\rho_{\mathrm{act}}^{\mathrm{free}}n_{\mathrm{adh}}\Delta t$ and $p_{\mathrm{myo}}^{\mathrm{on}}\approx k_{\mathrm{myo}}^{\mathrm{on}}\rho_{\mathrm{myo}}^{\mathrm{free}}n_{\mathrm{act}}\Delta t$, respectively, and the probabilities of an actin unit and myosin unit dissociating from a membrane/cortical element in an interval of time $\Delta t$ are, respectively, $p_{\mathrm{act}}^{\mathrm{off}}\approx k_{\mathrm{act}}^{\mathrm{off}} n_{\mathrm{act}}\Delta t$ and $p_{\mathrm{myo}}^{\mathrm{off}}\approx k_{\mathrm{myo}}^{\mathrm{off}} n_{\mathrm{myo}}\Delta t$. Actin and myosin at each cortical element are updated at each time step via random number generation, as described above. Old cortical nodes are only transiently retained in the spontaneous/stochastic blebbing model. Following a loss of adhesion between the membrane and the cortex, we assume that actin and myosin do not associate to the local detached cortex, and actin and myosin eventually disappear at the old cortex element after an imposed refractory time $t_{\mathrm{refr}}$.

The system of Eqs. (S1-S9) are computed in the Eulerian framework, whereas the forcing term that appear in Eq. (S3) come from the computation of membrane and cortex Lagrangian forces using Eqs. (S10) and (S16), respectively. To solve Eqs. (S1-S9), we thus make the following Lagrangian-Eulerian transformation

$$\mathcal{F}_{\mathrm{tot}}\left( \mathbf{x},t \right)=\int_{\Gamma_{m}} \mathbf{F}_{\mathbf{m}}\left( \xi,t \right)\delta\left( \mathbf{X}_{\mathbf{m}}\left( \xi,t \right)-\mathbf{x} \right)d\xi+\sum_{j=1}^{N_{b}} \delta\left( \mathbf{X}_{\mathbf{c}}\left( s_{j},t \right)-\mathbf{x} \right)\mathcal{F}_{\mathbf{c}}^{\mathbf{drag}}\left( \mathbf{x},t \right)-\mathbf{f}_{\mathbf{c, ecm}}\delta\left( \mathbf{X}_{\mathbf{ecm}}-\mathbf{x} \right) (S22)$$

where $\Gamma_{m}$ represents the membrane boundary. The transformation requires the discretization of the delta function in Eq. (S22). We use the well-behaved discretized delta function $\delta_{h}$ derived by Peskin^10^

$$\delta_{h}\left( \mathbf{x} \right)=\frac{1}{h^{2}}\phi\left( \frac{x_{1}}{h} \right)\phi\left( \frac{x_{2}}{h} \right) (S23)$$

$$\phi\left( r \right)=\left\{ \begin{aligned} 0 r\leq-2 \\ \frac{1}{8}\left( 5+2r-\sqrt{-7-12r-4r^{2}} \right) -2\leq r\leq-1 \\ \frac{1}{8}\left( 3+2r+\sqrt{1-4r-4r^{2}} \right) -1\leq r\leq0 \\ \frac{1}{8}\left( 3-2r+\sqrt{1+4r-4r^{2}} \right) 0\leq r\leq1 \\ \frac{1}{8}\left( 5-2r-\sqrt{-7+12r-4r^{2}} \right) 1\leq r\leq2 \\ 0 r\leq2 \end{aligned} \right. (S24)$$

Superposition of no-slip between the cell membrane and fluid and osmotic effects allows us to compute the membrane velocity $\mathbf{V}_{m}$ as follows

$$\mathbf{V}_{m}\left( s,t \right)=\int_{\Omega} \mathbf{v}_{f}\left( \mathbf{x},t \right)\delta\left( \mathbf{x-}\mathbf{X}_{\mathbf{m}}\left( s,t \right) \right)d\mathbf{x+}\zeta_{p}\left[ \left( \Delta\Pi\right)-\left( \Delta p \right) \right]\mathbf{n}_{\mathbf{m}} (S25)$$

where $\zeta_{p}$ is the membrane permeability coefficient, and $\Delta\Pi=h_{\mathrm{cell}}\mathrm{RT}\left( c_{\mathrm{in}}-c_{\mathrm{out}} \right)$ and $\Delta p=p_{\mathrm{in}}-p_{\mathrm{out}}$ are, respectively, the osmotic and hydrostatic pressure difference between the intracellular and extracellular domains. $h_{\mathrm{cell}}$ is the cell thickness in the z-direction, $R$ is the molar gas constant $(R=N_{A}k_{B}$, where $N_{A}$ is the Avogadro constant and $k_{B}$ is the Boltzmann constant), and $T$ is the absolute temperature, and we use the discretized delta function introduced in Eqs. (S23) and (S24) in Eq. (S25).

The intracellular and extracellular osmolyte concentrations, $c_{\mathrm{in}}$ and $c_{\mathrm{out}}$, obey the unsteady diffusion equation:

$$\frac{\partial c_{\mathrm{in}}}{\partial t}=\nabla\cdot\left( D_{\mathrm{in}}\nabla c_{\mathrm{in}} \right) \mathrm{in}\Omega_{\mathrm{in}}, \frac{\partial c_{\mathrm{out}}}{\partial t}=\nabla\cdot\left( D_{\mathrm{out}}\nabla c_{\mathrm{out}} \right) \mathrm{in}\Omega_{\mathrm{out}} (S26-S27)$$

where $D_{\mathrm{in}}$ and $D_{\mathrm{out}}$ are the intracellular and extracellular osmolyte diffusion coefficients, respectively. Although osmolyte advection can become important during bleb expansion, osmolyte transport is mainly driven throughout the whole bleb cycle dynamics by diffusion, thus we have neglected osmolyte advective transport in our model. Osmolyte transmembrane flux is facilitated by passive channels and active pumps. The flux continuity boundary conditions at the membrane reads:

$$D_{\mathrm{in}}\nabla c_{\mathrm{in}}\cdot\mathbf{n}_{\mathbf{m}}\boldsymbol{=}j_{\mathrm{pump}}+j_{\mathrm{passive}} \mathrm{on} \Gamma_{\mathrm{in}} (S28)$$

$$-D_{\mathrm{out}}\nabla c_{\mathrm{out}}\cdot\mathbf{n}_{\mathbf{m}}\boldsymbol{=}j_{\mathrm{pump}}+j_{\mathrm{passive}} \mathrm{on} \Gamma_{\mathrm{out}} (S29)$$

where $\Gamma_{\mathrm{in}}$ and $\Gamma_{\mathrm{out}}$ indicate that the boundary conditions are evaluated on the intracellular and extracellular sides on the membrane boundary, respectively. We assume that active pumps operate far from saturation conditions. Hence, the osmolyte flux from active pumping $j_{\mathrm{pump}}$ is a space- and time-dependent scalar $\alpha_{\mathrm{pump}}$, such that $j_{\mathrm{pump}}=\alpha_{\mathrm{pump}}\left( s,t \right)$. The osmolyte flux through passive channels is assumed to be of the form $j_{\mathrm{passive}}=\alpha_{\mathrm{passive}}\Delta c$ where we have neglected mechanosensitive effects. In most of our model results, we have taken the limit of infinitely rapid osmolyte diffusion ($D_{\mathrm{in}}\to\infty$, $D_{\mathrm{out}}\to\infty$), thus intracellular and extracellular osmolyte concentrations are spatially uniform. In this limiting case, cell area changes induced by variations in hydrostatic pressure modify the intracellular osmolyte concentration such that $c_{\mathrm{in}}\left( t \right)=c_{\mathrm{in}}\left( 0 \right)A_{\mathrm{cell}}(0)/A_{\mathrm{cell}}(t)$, where $A_{\mathrm{cell}}(t)$ is the cell area at time $t$. To solve the finite osmolyte diffusion coefficient problem, we use explicit Euler as the time integration scheme and the finite volume method^11^ to represent and evaluate the partial differential equation as an algebraic equation in two dimensions. This is numerically convenient, since the boundary conditions (S28) and (S29) are relatively easy to implement. For simplicity, we proceed to explain the numerical integration of Eq. (S26). A similar procedure has been used to integrate Eq. (S27). We divide the entire domain in equal-size square finite surfaces/cells and integrate Eq. (S26) over each cell $S(i,j)$:

$$\int_{S(i,j)} \frac{\partial c_{\mathrm{in}}}{\partial t}\mathrm{dS}=\int_{S(i,j)} \nabla\cdot\left( D_{\mathrm{in}}\nabla c_{\mathrm{in}} \right)\mathrm{dS} \mathrm{in}\Omega_{\mathrm{in}}, (S30)$$

where $i$ and $j$ indicate, respectively, the horizontal and vertical indices of the cell. We apply the divergence theorem on the right-hand-side, approximate integrals using the midpoint rule, and approximate the diffusive flux on each cell edge by using the second-order centered difference approximation provided that the all the neighbor cell centers lie in the intracellular space. Special treatment must be taken in cases where neighbor cell centers lie in the extracellular space. In these situations, one-sided centered difference approximations are utilized to approximate the diffusive flux across the cell edge. To enforce continuity of osmolyte flux across the cell boundary, we must estimate the osmolyte intracellular and extracellular concentration in points on the cell boundary. We estimate the osmolyte concentration on the cell boundary by applying bilinear interpolation making use of the osmolyte concentration of the nearest three cell centers.

The numerical solution at each time step is performed in six steps:

(I) membrane and cortical Lagrangian forces are computed from the membrane-cortex configurations using Eqs. (S10$-$S13) and (S15$-$S19)
(II) Total force on the viscoelastic fluid due to membrane and cortex is computed in the Eulerian framework using Eq. (S22), and fluid velocity and hydrostatic pressure are solved by applying the Fourier transform method to Eqs. (S1$-$S3)
(III) Fluid velocity at membrane and cortex Lagrangian nodes is computed, and membrane and cortex positions are updated using Eqs. (S25) and (S16) respectively
(IV) Polymer stress is updated by applying the Fourier transform to Eq. (S5) followed by the temporal integration of Eq. (S9)
(V) Number of membrane-cortex linkers engaged, cortical actin amounts, and cortical myosin amounts are updated by either integrating in time Eqs. (S14) and (S20$-$S21) when using the deterministic approach, or by using the stochastic model counterpart. In steps (IV) and (V), explicit Euler has been used as the time integration scheme
(VI) Osmolyte density is computed by solving the unsteady diffusion equations $(S26-S29)$, or by a simple calculation in the limit of infinitely fast osmolyte diffusion.

The forced/deterministic decohesion model was coded in MATLAB. Each simulated second required approximately four days to complete. Conversely, the spontaneous/stochastic blebbing model was coded in Fortran 90. We used resources from the Minnesota Supercomputing Institute (<https://msi.umn.edu>). Each spontaneous/stochastic simulation required approximately three days to complete.

We numerically solved for the fluid velocity $\mathbf{v}_{\mathbf{f}}$ and the hydrostatic pressure $p$ by using the fast Fourier transform. Let $\Psi_{k_{1}k_{2}}=\Psi\left( k_{1}h\mathbf{e}_{\mathbf{1}}+k_{2}h\mathbf{e}_{\mathbf{2}} \right)$, where $h=L/N$ is the grid size in both x and y directions, and $\mathbf{e}_{\mathbf{1}}$ and $\mathbf{e}_{\mathbf{2}}$ are the cartesian unit vectors. We define the discrete Fourier transformation of an arbitrary Eulerian function $\Psi$ as

$$\hat{\Psi}_{k_{1}k_{2}}=\frac{1}{N^{2}}\sum_{j_{1},j_{2}=0}^{N-1} e^{\left( {2\pi i}/N \right)\left( j_{1}k_{1}+j_{2}k_{2} \right)}\Psi(\mathbf{x}), 0\leq k_{1},k_{2} \leq N-1 (S31a)$$

where $k_{1}$ and $k_{2}$ represent the indices of the discrete Fourier transform (DFT) output in frequency domain. We define the inverse discrete Fourier transformation of $\Psi$ as

$$\Psi_{j_{1}j_{2}}=\sum_{k_{1},k_{2}=0}^{N-1} e^{-\left( {2\pi i}/N \right)\left( j_{1}k_{1}+j_{2}k_{2} \right)}\hat{\Psi}_{k_{1}k_{2}}=\sum_{k_{1},k_{2}=0}^{N-1} e^{-\left( {2\pi i}/L \right)\left( x_{1}k_{1}+x_{2}k_{2} \right)}\hat{\Psi}_{k_{1}k_{2}} (S31b)$$

We first apply the DFT to Eqs. (S1) and (S2):

$$\boldsymbol{-}\hat{\mathbf{D}}\hat{p}+\eta_{f}\hat{L}{\hat{\mathbf{v}}}_{\mathbf{f}}+\hat{\mathbf{D}}\boldsymbol{\cdot}{\hat{\boldsymbol{\sigma}}}_{\mathbf{p}}\boldsymbol{+}{\hat{\mathcal{F}}}_{\mathrm{tot}}=\boldsymbol{0} (S32)$$

$$\hat{\mathbf{D}}\boldsymbol{\cdot}{\hat{\mathbf{v}}}_{\mathbf{f}}\boldsymbol{=}0 (S33)$$

where the gradient/divergence $\hat{\mathbf{D}}$ and Laplacian $\hat{L}$ operators read

$${\hat{\mathbf{D}}}_{k_{1}k_{2}}=-\frac{i}{h}\sin\left( \frac{2\pi h}{L}\mathbf{k} \right)=\left( -\frac{i}{h}\sin\left( \frac{2\pi h}{L}k_{1} \right),-\frac{i}{h}\sin\left( \frac{2\pi h}{L}k_{2} \right) \right)$$

$$\hat{L}_{k_{1}k_{2}}=-\frac{4}{h^{2}}\sin\left( \frac{\pi}{N}\mathbf{k} \right)\cdot\sin\left( \frac{\pi}{N}\mathbf{k} \right)$$

Eliminating ${\hat{\mathbf{v}}}_{\mathbf{f}}$ by applying the divergence operator to Eq. (S32) allows us to derive an algebraic expression for the Fourier transform of the hydrostatic pressure:

$$\hat{p}_{k_{1}k_{2}}=\frac{\sin\left( \frac{2\pi}{N}\mathbf{k} \right)\cdot\sin\left( \frac{2\pi}{N}\mathbf{k} \right)\cdot{\hat{\boldsymbol{\sigma}}}_{\mathbf{p}}\boldsymbol{+}\mathrm{ih}\sin\left( \frac{2\pi}{N}\mathbf{k} \right)\cdot{\hat{\mathcal{F}}}_{\mathrm{tot}}}{\sin\left( \frac{2\pi}{N}\mathbf{k} \right)\cdot\sin\left( \frac{2\pi}{N}\mathbf{k} \right)} (S33)$$

Combining Eqs. (S32) and (S33) allows us to derive an algebraic expression for the Fourier transform of the fluid velocity:

$$\hat{v}_{k_{1}k_{2}}=\frac{1}{4\eta_{f}\sin\left( \frac{\pi}{N}\mathbf{k} \right)\cdot\sin\left( \frac{\pi}{N}\mathbf{k} \right)}\left[ -ih\sin\left( \frac{2\pi}{N}\mathbf{k} \right)\cdot{\hat{\boldsymbol{\sigma}}}_{\mathbf{p}}\mathbf{+}h^{2}{\hat{\mathcal{F}}}_{\mathrm{tot}}+ih\sin\left( \frac{2\pi}{N}\mathbf{k} \right)\frac{\sin\left( \frac{2\pi}{N}\mathbf{k} \right)\cdot\sin\left( \frac{2\pi}{N}\mathbf{k} \right)\cdot{\hat{\boldsymbol{\sigma}}}_{\mathbf{p}}\mathbf{+}\mathrm{ih}\sin\left( \frac{2\pi}{N}\mathbf{k} \right)\cdot{\hat{\mathcal{F}}}_{\mathrm{tot}}}{\sin\left( \frac{2\pi}{N}\mathbf{k} \right)\cdot\sin\left( \frac{2\pi}{N}\mathbf{k} \right)} \right] (S34)$$

Following a similar procedure, we have used the forward and inverse discrete Fourier transforms to compute the spatial gradients in Eq. (S9).

**Computation of the bleb nucleation correlation angle** $\mathbf{p}_{\mathbf{bleb}}$

We describe the location of each bleb by the polar angle between the first cortical node that mechanically dissociates from its corresponding plasma membrane node and a reference node, chosen arbitrarily, under the assumption that the cell maintains an approximately circular shape. We then define the bleb nucleation correlation angle as the polar angle difference between the position of two consecutive bleb nucleation events. This provides a practical measure of the angular separation between successive blebs.”

**Estimation of model parameters**

We proceed to estimate the different model parameters shown in Table S1.

**Table S1.** Model parameters

| Symbol | Description | Value | Legend/ References |
| --- | --- | --- | --- |
| $\mathbf{F}_{\mathbf{m}_{\mathbf{0}}}^{\mathbf{tens}}$ | Resting membrane tension | $2.5 pN\cdot{\mu m}^{-1}$ | 12 |
| $\boldsymbol{\kappa}_{\mathbf{m}}$ | Membrane spring stiffness | $\left[ 80-200 \right] pN\cdot{\mu m}^{-1}$ | 13,14 |
| ${\mathcal{l}_{\mathbf{m}}}_{\mathbf{0}}$ | Membrane spring resting length | $0.18 \mu m$ | A |
| $\boldsymbol{\beta}_{\mathbf{m}}$ | Effective two-dimensional membrane bending stiffness | $0.1 pN\cdot{\mu m}^{2}$ | 15 |
| ${\mathcal{l}_{\mathbf{adh}}}_{\mathbf{0}}$ | Resting length of membrane-cortex linkers | $0.5 \mu m$ | B |
| $\mathbf{k}_{\mathbf{adh}}^{\mathbf{on}}$ | Association rate constant of membrane-cortex linkers | $2K_{\tau} {\mu M}^{-1}\cdot s^{-1}$ | 16 |
| $\mathbf{k}_{\mathbf{adh}}^{\mathbf{off}}$ | Unloaded dissociation rate constant of membrane-cortex linkers | $0.2K_{\tau} s^{-1}$ | 17 |
| $\mathbf{F}_{\mathbf{adh}}^{\mathbf{rupt}}$ | Membrane-cortex linker rupture force | $\left[ 0.4-5 \right]$ $\mathrm{pN}$ | 18 |
| $\mathbf{N}_{\mathbf{adh}}^{\mathbf{tot}}$ | Total number of membrane-cortex linkers in the cell | $\left[ {10}^{5}-{10}^{6} \right]$ | Adjusted, C |
| $\boldsymbol{\kappa}_{\mathbf{adh}}$ | Effective stiffness of membrane-cortex linkers | $\left[ 0.5-5 \right] pN\cdot{\mu m}^{-1}$ | Adjusted |
| $\boldsymbol{\kappa}_{\mathbf{c}}$ | Cortex spring stiffness per actin unit | $\left[ 0.1-0.6 \right] pN\cdot{\mu m}^{-1}$ | Adjusted, D |
| ${\mathcal{l}_{\mathbf{c}}}_{\mathbf{0}}$ | Cortex spring resting length | $0.15 \mu m$ | A |
| $\mathbf{F}_{\mathbf{st}}$ | Myosin stall force per unit of actin and myosin | $1.9\times{10}^{-5}\mathrm{pN}$ | E |
| $\mathbf{v}_{\mathbf{0}}^{\mathbf{myo}}$ | Effective cortical unloaded myosin velocity | $\left[ 2-10 \right] \mu m\cdot s^{-1}$ | 19,20 |
| $\mathbf{k}_{\mathbf{act}}^{\mathbf{on}}$ | Actin association rate constant | $10K_{\tau} {\mu M}^{-1}\cdot s^{-1}$ | F |
| $\mathbf{k}_{\mathbf{act}}^{\mathbf{off}}$ | Actin dissociation rate constant | $0.3K_{\tau} s^{-1}$ | 17 |
| $\mathbf{k}_{\mathbf{myo}}^{\mathbf{on}}$ | Myosin association rate constant per unit of actin | $0.0025K_{\tau} {\mu M}^{-1}\cdot s^{-1}$ | G |
| $\mathbf{k}_{\mathbf{myo}}^{\mathbf{off}}$ | Myosin dissociation rate constant | $0.07K_{\tau} s^{-1}$ | 17,21 |
| $\mathbf{T}_{\mathbf{myo}}$ | Half of the period of cortical oscillations | $\left[ 2-12 \right]s$ | Adjusted, H |
| $\mathbf{K}_{\boldsymbol{\tau}}$ | Cortex turnover factor | $\left[ 1-200 \right]$ | I |
| $\mathbf{t}_{\mathbf{refr}}$ | Cortex refractory time | $50 ms$ | - |
| $\mathbf{N}_{\mathbf{act}}^{\mathbf{tot}}$ | Total number of actin units in the cell | $3.8\times{10}^{6}$ | J |
| $\mathbf{N}_{\mathbf{myo}}^{\mathbf{tot}}$ | Total number of myosin units in the cell | $\left[ 8\times{10}^{4}-2\times{10}^{5} \right]$ | Adjusted, K |
| $\boldsymbol{\gamma}_{\mathbf{c}}$ | Cortex drag coefficient | $[0.003-0.01] pN\cdot s\cdot{\mu m}^{-1}$ | Adjusted, L |
| $\boldsymbol{\xi}_{\mathbf{c}}$ | Mesh size cortex | $\left[ 0.015-0.2 \right]$ $\mu m$ | 22-24 |
| $\mathbf{r}_{\mathbf{c}}$ | Effective actin filament radius | $3.5 nm$ | 25 |
| $\boldsymbol{\eta}_{\mathbf{f}}$ | Effective two-dimensional dynamic viscosity of fluid component | $\left[ 0.1-1.4 \right] pN\cdot s\cdot{\mu m}^{-1}$ | M |
| $\boldsymbol{\eta}_{\mathbf{p}}^{\mathbf{in}}$ | Intracellular polymer viscosity | $\left[ 0.1-10 \right] pN\cdot s\cdot{\mu m}^{-1}$ | Adjusted, N |
| $\boldsymbol{\eta}_{\mathbf{p}}^{\mathbf{out}}$ | Extracellular polymer viscosity | $\left[ 0.1-10 \right] pN\cdot s\cdot{\mu m}^{-1}$ | Adjusted, N |
| $\boldsymbol{\lambda}_{\mathbf{p}}^{\mathbf{in}}$ | Intracellular polymer relaxation time | $\left[ {10}^{-2}-{10}^{2} \right] s$ | Adjusted, N |
| $\boldsymbol{\lambda}_{\mathbf{p}}^{\mathbf{out}}$ | Extracellular polymer relaxation time | $\left[ {10}^{-2}-{10}^{2} \right] s$ | Adjusted, N |
| $\mathbf{L}_{\mathbf{p}}$ | Polymer extensibility parameter | $10$ | O |
| $\boldsymbol{\zeta}_{\mathbf{p}}$ | Membrane permeability coefficient | ${2.8\times10}^{-4} {\mu m}^{2}\cdot\mathrm{pN}^{-1}\cdot s^{-1}$ | P |
| $\mathbf{c}_{\mathbf{in}}\mathbf{(0)}$ | Initial intracellular osmolyte concentration | $339.5283 mM$ | 26 |
| $\mathbf{c}_{\mathbf{out}}\mathbf{(0)}$ | Initial extracellular osmolyte concentration | $339.5 mM$ | 26 |
| $\mathbf{D}_{\mathbf{in}}$ | Intracellular osmolyte diffusion coefficient | $27 {\mu m}^{2}\cdot s^{-1}$ | 27 |
| $\mathbf{D}_{\mathbf{out}}$ | Extracellular osmolyte diffusion coefficient | $\left[ 30-2000 \right]$ ${\mu m}^{2}\cdot s^{-1}$ | 28 |
| $\boldsymbol{\kappa}_{\mathbf{cell-ecm}}$ | Effective stiffness of the cell adhesion protein complex-extracellular matrix tandem | $\left[ 0-500 \right]$ $\mathrm{pN}\cdot{\mu m}^{-1}$ | Adjusted, Q |
| $\mathbf{R}_{\mathbf{cell}}\boldsymbol{(0)}$ | Initial cell radius | $3 \mu m$ | O |
| $\mathbf{h}_{\mathbf{cell}}$ | Cell thickness in z-direction | $6 \mu m$ | O |
| $\mathbf{N}_{\mathbf{b}}$ | Number of membrane and cortex discretization nodes | $100$ | O |
| $\mathbf{N}$ | Number of Eulerian nodes used to discretize the whole domain $\Omega$ | $256$ | - |
| $\mathbf{L}$ | Domain size in x and y directions | $50$ $\mu m$ | - |
| $\boldsymbol{\phi}_{\mathbf{b}}$ | Fraction of the plasma membrane perimeter that loses mechanical connection with the underlying cortex in the deterministic model | $0.15$ | R |
| $\boldsymbol{\Delta t}$ | Time step in all simulations | ${10}^{-6}s$ | - |

**A.** Computed at the onset of our simulations as the separation between neighbor membrane Lagrangian nodes ${\mathcal{l}_{m}}_{0}=2\pi R_{\mathrm{cell}}(0)/N_{b}$ and neighbor cortical nodes ${\mathcal{l}_{c}}_{0}=2\pi\left( R_{\mathrm{cell}}\left( 0 \right)-h_{c} \right)/N_{b}$.

**B.** The mean cortex thickness is $\approx230 \mathrm{nm}$ in a Jurkat cell line^29^. We choose ${\mathcal{l}_{\mathrm{adh}}}_{0}\boldsymbol{=}0.5 \mu m$, a larger membrane-cortex separation to reduce the required spatial resolution of the Eulerian grid and associated computational cost.

**C.** We assume that the mean number of membrane-cortex linkers at each cortex node $\bar{n}_{\mathrm{adh}}$ is approximately equal to the mean number of actin filaments $\bar{n}_{\mathrm{adh}}=36$ (refer to F). We can then estimate the cytoplasmic density of membrane-cortex linkers from a kinetic balance of linkers in the cortex as $\rho_{\mathrm{adh}}^{\mathrm{free}}=\left( {k_{\mathrm{adh}}^{\mathrm{off}}}/{k_{\mathrm{adh}}^{\mathrm{on}}} \right)\bar{n}_{\mathrm{adh}}=3.6\mu M$. The total number of membrane-cortex linkers in the cell can then be obtained as $N_{\mathrm{adh}}^{\mathrm{tot}}=\rho_{\mathrm{adh}}^{\mathrm{free}}A_{\mathrm{cell}}(0)h_{\mathrm{cell}}+N_{b}\bar{n}_{\mathrm{adh}}\approx3.8\times{10}^{5}$.

**D.** From the effective cortex stiffness coefficient previously reported $E_{c}\approx{10}^{4} pN\cdot{\mu m}^{-1}$ ^30^, we can estimate the cortex spring stiffness per actin unit as: $\kappa_{c}={E_{c}}/{\bar{n}_{\mathrm{act}}}\approx0.5 pN\cdot{\mu m}^{-1}$, where $\bar{n}_{\mathrm{act}}$ is the estimated mean number of actin units at each cortical node (refer to legend J).

**E.** We estimate the myosin stall force per unit of actin and myosin from the reported values of cortical tension, which lie in the range $T_{c}\approx\left[ 55-1600 \right] pN\cdot{\mu m}^{-1}$ ^30-32^. Assuming a cortical tension of $T_{c}=200 \mathrm{pN}$, then $F_{\mathrm{st}}={T_{c}}/{\bar{n}_{\mathrm{myo}}\bar{n}_{\mathrm{act}}}=1.9\times{10}^{-5} \mathrm{pN}$. Here, $\bar{n}_{\mathrm{myo}}$ is the mean number of myosin units at each cortical node. The chosen cortical tension is on the lower end of reported values. However, we study a wide range of cortical tensions by varying the total number of myosin molecules in the cell, such that the range of experimental values is encompassed by our simulations.

**F.** The actin association rate constant for each actin filament barbed end is $k_{\mathrm{barbed}}^{\mathrm{on}}\approx10 {\mu M}^{-1}\cdot s^{-1}$ ^33,34^. Assuming that the mean number of actin filaments per unit length in our two-dimensional cortex is approximately of the same order to the well characterized fission yeast cytokinetic ring $\rho_{\mathrm{fil}}\approx31.8 fil/\mu m$^35^, then the mean number of actin filaments at each cortical node is $\bar{n}_{\mathrm{fil}}=2\pi R_{\mathrm{cell}}h_{\mathrm{cell}}\rho_{\mathrm{fil}}/N_{b}\approx36 fil$. The actin association rate at each cortical node is then $k_{\mathrm{act}}^{\mathrm{on}}={\bar{n}_{\mathrm{fil}}k_{\mathrm{barbed}}^{\mathrm{on}}}/{\bar{n}_{\mathrm{adh}}}=10 {\mu M}^{-1}\cdot s^{-1}.$

**G.** We assume that the number of myosin molecules per unit length in our two-dimensional cortex is approximately that in the fission yeast cytokinetic ring $\rho_{\mathrm{myo}}\approx455 molecules/\mu m$^35^. Then, the mean number of myosin polypeptides at each cortical node is $\bar{n}_{\mathrm{myo}}=2\pi R_{\mathrm{cell}}h_{\mathrm{cell}}\rho_{\mathrm{myo}}/N_{b}\approx515$. Assuming that the concentration of cytoplasmic myosin is $\rho_{\mathrm{myo}}^{\mathrm{free}}=0.7\mu M$ (refer to I), then we can estimate the myosin association rate constant per unit of actin from a kinetic balance of myosin in the cortex: $k_{\mathrm{myo}}^{\mathrm{on}}={k_{\mathrm{myo}}^{\mathrm{off}}\bar{n}_{\mathrm{myo}}}/{\rho_{\mathrm{myo}}^{\mathrm{free}}\bar{n}_{\mathrm{act}}}\approx0.0025 {\mu M}^{-1}\cdot s^{-1}$.

**H.** The range of the period of oscillations has been chosen such that a few oscillatory cycles are simulated within the total simulation time (usually 18 seconds).

**I.** We introduce a cortex turnover factor to accelerate the kinetics of membrane-cortex linkers, actin and myosin, allowing us to observe many bleb cycles in the total simulated time, thus increasing acquired data to achieve statistical significance.

**J.** The amount of actin molecules per unit length in the fission yeast ring is $\rho_{\mathrm{act}}\sim1.8\times{10}^{4}$ $molecules/\mu m$^35^. We want to estimate an upper limit for $N_{\mathrm{act}}^{\mathrm{tot}}$. We then assume that our two-dimensional cellular cortex contains a similar actin density to that of the fission yeast ring, then the number mean number of F-actin monomers at each cortical node in our model is $\bar{n}_{\mathrm{act}}={2\pi R_{\mathrm{cell}}h_{\mathrm{cell}}\rho_{\mathrm{act}}}/{N_{b}}\approx2.04\times{10}^{4}$. Notice that we have scaled the number of actin units in each node by the cell thickness. Using the reported values of actin association and dissociation rate constants and the number of actin units at each cortical node, we can estimate the mean concentration of G-actin units in the cytoplasm from Eq. (12): $\rho_{\mathrm{act}}^{\mathrm{free}}={\bar{n}_{\mathrm{act}}k_{\mathrm{act}}^{\mathrm{off}}}/{k_{\mathrm{act}}^{\mathrm{on}}}=17\mu M$, which is largely equal to half the intracellular G-actin concentration ($30-37$ $\mu M$) that have been reported in the literature^36^. The total number of actin units in the cell is then: $N_{\mathrm{act}}^{\mathrm{tot}}=\rho_{\mathrm{act}}^{\mathrm{free}}A_{\mathrm{cell}}(0)h_{\mathrm{cell}}+N_{b}\bar{n}_{\mathrm{act}}\approx3.8\times{10}^{6}$.

**K.** The cytoplasmic concentration of the different nonmuscle myosin II isoforms (NMIIA, NMIIB, NMIIC) has been measured in HeLa cells and a few pancreatic cancer cell lines^37^. According to these measurements, the total free myosin concentration is within the range $\rho_{\mathrm{myo}}^{\mathrm{free}}\sim\left[ 0.6-0.8 \right] \mu M$. Then the total number of myosin II molecules in the cell can be estimated as $N_{\mathrm{myo}}^{\mathrm{tot}}=\rho_{\mathrm{myo}}^{\mathrm{free}}A_{\mathrm{cell}}(0)h_{\mathrm{cell}}+N_{b}\bar{n}_{\mathrm{myo}}$. We find that $N_{\mathrm{myo}}^{\mathrm{tot}}$is then within the range $\left[ 1.13\times{10}^{5}-1.33\times{10}^{5} \right]$. Notice that we have used the value of $\bar{n}_{\mathrm{myo}}$ estimated in G. We therefore adjust $N_{\mathrm{myo}}^{\mathrm{tot}}$ within a reasonable range $\left[ 8\times{10}^{4}-2\times{10}^{5} \right]$.

**L.** We estimate the drag coefficient experienced by cytoplasmic material as it goes through the cortex by considering the transverse flow of a Newtonian fluid through an array of infinite parallel rods. Assuming a reasonable cortical mesh size $\xi_{c}\approx0.015 \mu m$^22^, the volume fraction of actin filaments in the cortex is $\phi_{c}={\pi r_{c}^{2}}/{\xi_{c}^{2}}\approx0.1710$. The drag coefficient $\gamma_{c}^{'}$ per unit length can then be obtained by solving the following non-linear algebraic equation^38^:

$$\frac{4\pi\eta_{f}^{3D}}{\gamma_{c}^{'}}+\ln\left( \sqrt{\frac{\gamma_{c}^{'}}{4\pi\eta_{f}^{3D}}} \right)+\gamma_{\mathrm{eul}}-0.47\frac{\gamma_{c}^{'}}{4\pi\eta_{f}^{3D}}+\ln\left( \sqrt{\phi_{c}} \right)=0$$

where $\eta_{f}^{3D}$ is the cytoplasmic viscosity, and $\gamma_{\mathrm{eul}}$ is the Euler-Mascheroni constant. Using a cytoplasmic viscosity of $\eta_{f}^{3D}=0.005 pN\cdot s\cdot{\mu m}^{-2}$, we get $\gamma_{c}^{'}\approx0.016 pN\cdot s\cdot{\mu m}^{-2}$. The drag coefficient associated to each cortical node can then be obtained by multiplying $\gamma_{c}^{'}$ by the characteristic distance between cortical nodes ${\mathcal{l}_{c}}_{0}$. We get: $\gamma_{c}\approx0.003 pN\cdot s\cdot{\mu m}^{-1}$.

**M.** Typical values of cytoplasmic viscosity are $\left[ {2\times10}^{-3}-10 \right] pN\cdot s\cdot{\mu m}^{-2}$ ^39-41^. We estimate the effective two-dimensional cytoplasmic viscosity as $\eta_{f}\boldsymbol{=}h_{\mathrm{cell}}\eta_{f}^{3D}\approx\left[ 0.01-60 \right] pN\cdot s\cdot{\mu m}^{-1}$. We use a wider range of fluid viscosity values to study statistically significant differences in cell migratory behavior.

**N.** We explore how different mechanical properties of the cytoplasm and extracellular space influence cell migration capabilities. To explore maximal theoretical migration speeds achieved by bleb-producing cells, we use the viscous fluid limit in most of our simulations.

**O.** Arbitrarily chosen.

**P.** The membrane permeability coefficient $\zeta_{p}$ used in Eq. (S25) can be written as $\zeta_{p}=\upsilon_{0}/(k_{B}Tc_{w})$, where $\upsilon_{0}$ is the osmotic water permeability commonly estimated in experiments $\upsilon_{0}\sim\left[ 5-100 \right] \mu m\cdot s^{-1}$ ^42^, and $c_{w}^{-1}$ is the volume of a water molecule. Our model is two-dimensional; thus, we will use the corresponding cross-sectional area of a water molecule $A_{w}$ instead. The effective radius of a water molecule can be estimated from $c_{w}\approx55.55 mol\cdot L^{-1}$, we get $R_{w}\approx1.9\times{10}^{-4} \mu m$, and $A_{w}\approx1.16\times{10}^{-7}{\mu m}^{2}$. We choose $\upsilon_{0}=10 \mu m\cdot s^{-1}$, thus $\zeta_{p}=\upsilon_{0}A_{w}/(k_{B}T)\approx{2.8\times10}^{-4} {\mu m}^{2}\cdot\mathrm{pN}^{-1}\cdot s^{-1}$.

**Q.** To study the migratory potential of adherent blebby cells (Fig. 6), we set $\kappa_{cell-ecm}=500 pN\cdot{\mu m}^{-1}$, a stiff enough cellular adhesion protein complex-extracellular matrix that effectively resists rearward cortical forces during bleb retraction.

**R.** Prescribed fraction of the plasma membrane perimeter (neck bleb size) that loses mechanical connection with the underlying cortex used in Figs. 2, 6, S1A, S1B, S2 and S6. The number of membrane-cortex bonds broken is 15.

**Table S2. Parameter values used to produce the model results, unless otherwise specified**

| Symbol | Value |
| --- | --- |
| $\mathbf{F}_{\mathbf{m}_{\mathbf{0}}}^{\mathbf{tens}}$ | $2.5 pN\cdot{\mu m}^{-1}$ |
| $\boldsymbol{\kappa}_{\mathbf{m}}$ | $120 pN\cdot{\mu m}^{-1}$ |
| ${\mathcal{l}_{\mathbf{m}}}_{\mathbf{0}}$ | $0.18 \mu m$ |
| $\boldsymbol{\beta}_{\mathbf{m}}$ | $0 pN\cdot{\mu m}^{2}$ |
| ${\mathcal{l}_{\mathbf{adh}}}_{\mathbf{0}}$ | $0.5 \mu m$ |
| $\mathbf{k}_{\mathbf{adh}}^{\mathbf{on}}$ | $20K_{\tau} {\mu M}^{-1}\cdot s^{-1}$ |
| $\mathbf{k}_{\mathbf{adh}}^{\mathbf{off}}$ | $0.4K_{\tau} s^{-1}$ |
| $\mathbf{F}_{\mathbf{adh}}^{\mathbf{rupt}}$ | $0.4\mathrm{pN}$ |
| $\mathbf{N}_{\mathbf{adh}}^{\mathbf{tot}}$ | $3.5\times{10}^{5}$ |
| $\boldsymbol{\kappa}_{\mathbf{adh}}$ | $5pN\cdot{\mu m}^{-1}$ |
| $\boldsymbol{\kappa}_{\mathbf{c}}$ | $0.1 pN\cdot{\mu m}^{-1}$ |
| ${\mathcal{l}_{\mathbf{c}}}_{\mathbf{0}}$ | $0.15 \mu m$ |
| $\mathbf{F}_{\mathbf{st}}$ | $1.9\times{10}^{-5}\mathrm{pN}$ |
| $\mathbf{v}_{\mathbf{0}}^{\mathbf{myo}}$ | $10 \mu m\cdot s^{-1}$ |
| $\mathbf{k}_{\mathbf{act}}^{\mathbf{on}}$ | $10K_{\tau} {\mu M}^{-1}\cdot s^{-1}$ |
| $\mathbf{k}_{\mathbf{act}}^{\mathbf{off}}$ | $0.3K_{\tau} s^{-1}$ |
| $\mathbf{k}_{\mathbf{myo}}^{\mathbf{on}}$ | $0.00125K_{\tau} {\mu M}^{-1}\cdot s^{-1}$ |
| $\mathbf{k}_{\mathbf{myo}}^{\mathbf{off}}$ | $0.035K_{\tau} s^{-1}$ |
| $\mathbf{T}_{\mathbf{myo}}$ | $4s$ |
| $\mathbf{K}_{\boldsymbol{\tau}}$ | $1$ |
| $\mathbf{t}_{\mathbf{refr}}$ | $50 ms$ |
| $\mathbf{N}_{\mathbf{act}}^{\mathbf{tot}}$ | $3.8\times{10}^{6}$ |
| $\mathbf{N}_{\mathbf{myo}}^{\mathbf{tot}}$ | $1.4\times{10}^{5}$ |
| $\boldsymbol{\gamma}_{\mathbf{c}}$ | $0.003 pN\cdot s\cdot{\mu m}^{-1}$ |
| $\boldsymbol{\xi}_{\mathbf{c}}$ | $0.015 \mu m$ |
| $\mathbf{r}_{\mathbf{c}}$ | $3.5 nm$ |
| $\boldsymbol{\eta}_{\mathbf{f}}$ | $0.4 pN\cdot s\cdot{\mu m}^{-1}$ |
| $\boldsymbol{\eta}_{\mathbf{p}}^{\mathbf{in}}$ | $0 pN\cdot s\cdot{\mu m}^{-1}$ |
| $\boldsymbol{\eta}_{\mathbf{p}}^{\mathbf{out}}$ | $0 pN\cdot s\cdot{\mu m}^{-1}$ |
| $\boldsymbol{\lambda}_{\mathbf{p}}^{\mathbf{in}}$ | $\infty$ |
| $\boldsymbol{\lambda}_{\mathbf{p}}^{\mathbf{out}}$ | $\infty$ |
| $\mathbf{L}_{\mathbf{p}}$ | $10$ |
| $\boldsymbol{\zeta}_{\mathbf{p}}$ | ${2.8\times10}^{-4} {\mu m}^{2}\cdot\mathrm{pN}^{-1}\cdot s^{-1}$ |
| $\mathbf{c}_{\mathbf{in}}\mathbf{(0)}$ | $339.5283 mM$ |
| $\mathbf{c}_{\mathbf{out}}\mathbf{(0)}$ | $339.5 mM$ |
| $\mathbf{D}_{\mathbf{in}}$ | $\infty$ |
| $\mathbf{D}_{\mathbf{out}}$ | $\infty$ |
| $\boldsymbol{\kappa}_{\mathbf{cell-ecm}}$ | $0$ $\mathrm{pN}\cdot{\mu m}^{-1}$ |
| $\mathbf{R}_{\mathbf{cell}}\boldsymbol{(0)}$ | $3 \mu m$ |
| $\mathbf{h}_{\mathbf{cell}}$ | $6 \mu m$ |
| $\mathbf{N}_{\mathbf{b}}$ | $100$ |
| $\boldsymbol{\phi}_{\mathbf{b}}$ | $0.15$ |

11.	Moukalled, F., Mangani, L., Darwish, M., and SpringerLink. (2016). The Finite Volume Method in Computational Fluid Dynamics : An Advanced Introduction with OpenFOAM® and Matlab, 1st 2015. Edition (Springer International Publishing : Imprint: Springer).

12.	Dai, J., and Sheetz, M.P. (1999). Membrane tether formation from blebbing cells. Biophys J *77*, 3363-3370. 10.1016/S0006-3495(99)77168-7.

13.	Hochmuth, R.M. (2000). Micropipette aspiration of living cells. J Biomech *33*, 15-22. 10.1016/s0021-9290(99)00175-x.

14.	Needham, D., and Hochmuth, R.M. (1992). A sensitive measure of surface stress in the resting neutrophil. Biophys J *61*, 1664-1670. 10.1016/S0006-3495(92)81970-7.

15.	Khelashvili, G., Kollmitzer, B., Heftberger, P., Pabst, G., and Harries, D. (2013). Calculating the Bending Modulus for Multicomponent Lipid Membranes in Different Thermodynamic Phases. J Chem Theory Comput *9*, 3866-3871. 10.1021/ct400492e.

16.	Korkmazhan, E., and Dunn, A.R. (2022). The membrane-actin linker ezrin acts as a sliding anchor. Sci Adv *8*, eabo2779. 10.1126/sciadv.abo2779.

25.	Cooper, G.M., Hausman, R.E., and Hausman, R.E. (2007). The cell: a molecular approach (ASM press).

32.	Fischer-Friedrich, E., Hyman, A.A., Jülicher, F., Müller, D.J., and Helenius, J. (2014). Quantification of surface tension and internal pressure generated by single mitotic cells. Sci Rep *4*, 6213. 10.1038/srep06213.

33.	Pollard, T.D. (2016). Actin and Actin-Binding Proteins. Cold Spring Harb Perspect Biol *8*. 10.1101/cshperspect.a018226.
